# Comparing serial X-ray crystallography and microcrystal electron diffraction (MicroED) as methods for routine structure determination from small macromolecular crystals

**DOI:** 10.1101/767061

**Authors:** Alexander M Wolff, Iris D Young, Raymond G Sierra, Aaron S Brewster, Michael W Martynowycz, Eriko Nango, Michihiro Sugahara, Takanori Nakane, Kazutaka Ito, Andrew Aquila, Asmit Bhowmick, Justin T Biel, Sergio Carbajo, Aina E Cohen, Saul Cortez, Ana Gonzalez, Tomoya Hino, Dohyun Im, Jake D Koralek, Minoru Kubo, Tomas S Lazarou, Takashi Nomura, Shigeki Owada, Avi Samelson, Rie Tanaka, Tomoyuki Tanaka, Erin M Thompson, Henry van den Bedem, Rahel A Woldeyes, Fumiaki Yumoto, Wei Zhao, Kensuke Tono, Sébastien Boutet, So Iwata, Tamir Gonen, Nicholas K Sauter, James S Fraser, Michael C Thompson

**Author notes:** MRC laboratory of molecular biology, Cambridge, UK. **Synopsis** The authors perform serial X-ray crystallography and microcrystal electron diffraction (MicroED) using microcrystals of the enzyme cyclophilin A. Their results highlight the strengths and weakness of the two complementary methods.

## Abstract

Innovative new crystallographic methods are facilitating structural studies from ever smaller crystals of biological macromolecules. In particular, serial X-ray crystallography and microcrystal electron diffraction (MicroED) have emerged as useful methods for obtaining structural information from crystals on the nanometer to micron scale. Despite the utility of these methods, their implementation can often be difficult, as they present many challenges not encountered in traditional macromolecular crystallography experiments. Here, we describe XFEL serial crystallography experiments and MicroED experiments using batch-grown microcrystals of the enzyme cyclophilin A (CypA). Our results provide a roadmap for researchers hoping to design macromolecular microcrystallography experiments, and they highlight the strengths and weaknesses of the two methods. Specifically, we focus on how the different physical conditions imposed by the sample preparation and delivery methods required for each type of experiment effect the crystal structure of the enzyme.

## 1. Introduction

In macromolecular crystallography, collecting full datasets from small crystals has been a challenge, because their weaker diffracting power limits the amount of signal that can be successfully obtained before the effects of X-ray radiation damage become significant. Thus, crystallographers have always been faced with a practical need to either optimize the growth of relatively large crystals, or to make the most of smaller crystals by implementing clever data collection and merging strategies (Cusack *et al.*, 1998; Zander *et al.*, 2015). Methodological advances that facilitated measurement of diffraction data from smaller and smaller crystals, such as the introduction of crystal cryocooling and the development of microfocus X-ray beams, have enabled structure determination of increasingly challenging targets, for which large crystals could never be obtained (Liu *et al.*, 2013; Zhou *et al.*, 2016). Additionally, small crystals have proven advantageous in a number of other contexts. For example, when crystals display long-range disorder, such as high mosaicity, a reduction in the total number of mosaic blocks reduces the spread of Bragg peaks and generally improves the overall data quality (Chernov, 1999). Small crystals are also advantageous when the diffraction experiment is preceded by a perturbation to the crystal. This includes common crystal treatments such as flash cooling or ligand soaking, as well as more uncommon perturbations, like stimulation of crystallized molecules for time-resolved experiments (Coquelle *et al.*, 2018; Olmos *et al.*, 2018). Because they have substantially less volume and a limited number of unit cells, perturbations can be applied more rapidly and uniformly to smaller crystals than to larger ones, and smaller crystals accumulate less stress resulting from changes in crystal lattice dimensions. The development of protein “microcrystallography” techniques, which are optimized for measuring crystals whose physical dimensions are tens of microns or smaller, has offered access to these opportunities and benefits, and led to a shift in what is considered to be a valuable specimen for experimental characterization.

The past decade has seen an explosion of new technology for protein microcrystallography. The increased brightness available for crystallography at modern X-ray light sources, including synchrotrons and X-ray free-electron lasers (XFELs), led to the development of “serial crystallography.” In a serial crystallography experiment, the X-ray beam is typically very bright and tightly focused, so that extremely short exposure times produce measurable diffraction images, even for very small crystals (Chapman *et al.*, 2011). The intense X-ray beams used for these experiments destroy the samples rapidly, allowing only a single diffraction image to be collected per crystal. Therefore the crystals must be rapidly (or “serially”) replenished at the X-ray interaction point in order for the experiment to be efficient. By measuring single diffraction snapshots of many randomly-oriented crystals, it is possible to completely sample the reflections in reciprocal space and integrate the Bragg intensities. Importantly, serial crystallography experiments are generally conducted at room temperature, potentially giving more physiologically-relevant insight into molecular structure by avoiding the artifacts associated with cryocooling.

Alongside the development of serial crystallography, major recent breakthroughs have been made in the field of microcrystal electron diffraction (MicroED). Specifically, the facile collection of continuous rotation datasets (Nannenga *et al.*, 2014; Shi *et al.*, 2013) from flash-cooled microcrystals using a standard transmission electron microscope operated in diffraction mode. Because the microscopes required for MicroED are now widespread as a result of booming interest in electron cryomicroscopy (cryoEM), MicroED holds great potential for the determination of both protein and small molecule structures (Nannenga & Gonen, 2019). Collectively, these new frontiers in macromolecular microcrystallography have created new opportunities for structural biology. Examples include structure determination from crystals as small as a few hundred nanometers in each of their dimensions (Chapman *et al.*, 2011; Nannenga & Gonen, 2019; de la Cruz *et al.*, 2017), and a new generation of challenging time-resolved measurements (Young *et al.*, 2016; Nango *et al.*, 2016) at high spatial and temporal resolution.

Despite the interesting possibilities that are now within reach, the optimization of sample preparation and data collection protocols for microcrystallography experiments remains challenging. First, it is necessary to decide which measurement technique (i.e. X-ray or MicroED) is best suited to a given sample or research question. Then, if appropriately-sized crystals are not obtained serendipitously, the experimenter must generate microcrystals with the correct size and density (crystals/uL), either by targeted growth or by manipulation of larger crystal specimens. Next, it is essential to choose an appropriate method for delivering the microcrystals to the X-ray beam or to the column of the electron microscope. In the case of serial X-ray crystallography, multiple strategies have been explored for rapidly replenishing crystals at the X-ray interaction point. In addition to fixed-target approaches (Baxter *et al.*, 2016; Fuller *et al.*, 2017; Hunter *et al.*, 2014), where crystals are mounted on a solid support and moved through the X-ray interaction region using automation, several methods have come into widespread use that exploit microfluidics to create free-standing streams, or “jets,” of microcrystal slurries. Various different types of microfluidic devices, collectively referred to as “sample injectors,” have been developed for this purpose *(Sierra et al., 2012a; Weierstall et al., 2014).* Each type uses a different physical principle for carrying microcrystals to the X-ray beam by generating a stream of flowing liquid that is tens to hundreds of microns in diameter, and each method subjects the crystals to different conditions which could potentially affect the quality of data acquired or the structure of the molecule itself. These conditions include exposure to strong electric fields (Sierra *et al.*, 2012*a*), high pressure (Weierstall *et al.*, 2014), and additives (Sugahara *et al.*, 2017) that change the chemical properties of mother liquor solutions.

Similarly in MicroED several sample preparation methods have been reported such as direct pipetting of nanocrystals onto EM grids (Rodriguez *et al.*, 2015); sonication, vortexing, vigorous pipetting, or crushing to break big crystals into fragments and create a nanocrystal slurry (de la Cruz *et al.*, 2017). Crystals are then drop cast onto EM grids, traditionally used for cryoEM, then the grids are blotted of excess solvent and flash frozen in supercooled ethane. Once frozen, microcrystals prepared in this fashion can be used directly for data collection, or they can be subjected to a milling procedure that utilizes a scanning electron microscope with a focused ion beam (FIB-SEM) to create crystalline lamellae of the desired thickness (Martynowycz *et al.*, 2019*a*,*b*). Because electrons interact with matter more strongly than X-rays do, the ideal crystal thickness for MicroED measurements is only several hundred nanometers (Martynowycz *et al.*, 2019*b*) and the milling process is critical for samples that exceed this thickness. As for serial crystallography, sample preparation and delivery for MicroED involves subjecting crystals to unusual conditions that are not typically encountered when samples are prepared for traditional crystallographic experiments. These conditions include dehydration and exposure to shear forces that are produced by the flow of solvent during blotting (Martynowycz *et al.*, 2019*a*), as well as potential damage induced by FIB milling. All of these considerations create a complex landscape, and designing the best experiment for a new system of interest is often a non-trivial process. This calculus is further complicated by the fact that the extent to which the unusual experimental conditions affect the quality of data acquired, or the structure of the molecule itself, have not been rigorously characterized.

Here, we discuss the planning, optimizing, and executing protein microcrystallography experiments, using human cyclophilin A (CypA) as a model protein system. CypA is a proline isomerase enzyme that is highly abundant in human cells, and plays important biological roles as both a protein folding chaperone and modulator of intracellular signaling pathways. Prior work has shown that CypA readily forms large crystals, which have been successfully used for traditional rotation crystallography at synchrotron X-ray sources and for fixed target measurements at an XFEL source (Fraser *et al.*, 2009; Keedy *et al.*, 2015). Starting from crystallization conditions that produce large (hundreds of microns in each dimension) CypA crystals, we optimized the preparation of high-density microcrystal slurries. We then used these samples for a variety of microcrystallography experiments, including serial X-ray crystallography with three different microfluidic sample injectors, and MicroED. Because the data collected across the different types of experiments were derived from similarly prepared microcrystal samples, and analyzed using the same protocols, we were able to perform a rigorous comparison of the results. For each method, we evaluate the robustness of sample preparation and delivery, the statistical quality of the measured data, and the properties of the resulting atomic models. Our results illustrate the inherent strengths and weaknesses of these new and exciting techniques for macromolecular microcrystallography, and lay out a roadmap for optimization of this promising category of experiments.

## 2. Methods

### 2.1 Protein Expression & Purification

Wild-type human Cyclophilin A (CypA) was expressed and purified as previously described (Fraser *et al.*, 2009). Briefly, following purification, the protein was stored in a solution of 20 mM HEPES pH 7.5, 20 mM NaCl, 0.5 mM TCEP at 4° C until use. Finally, samples were concentrated using amicon centrifugal filters, then crystallized as described below.

### 2.2 Crystal Formation & Optimization

For exploration of CypA’s crystallization phase space, crystallization trays were set as follows. Well solutions containing 100 mM HEPES pH 7.5, 5 mM TCEP, and PEG 3350 (concentration varied) were distributed into Greiner 96-well Imp@ct microbatch crystallization plates. Each well contained 2 μL of the respective well solution, mixed with 2 μL of protein at the respective concentration. These drops were then vapor sealed using 12 μL of paraffin oil. For large-scale production of crystals in batch, 600 μL of protein at 60 mg/mL was combined with 400 μL of 50% PEG 3350 in a glass vial and stirred with a stir bar at a constant rate (RPM varied). Crystallization was robust over a temperature range spanning 20-25°C.

### 2.3 Crystal Analysis

Raw images of microcrystal slurries were taken under a Nikon Ti microscope in differential interference contrast mode, with a Nikon DS-Qi2 camera. Data were interpreted using Fiji software (Rueden *et al.*, 2017). In addition to imaging crystalline slurries, particle densities (crystals/mL) were analyzed using an INCYTO C-Chip. Diffraction tests were carried out at Stanford Synchrotron Radiation Lightsource (SSRL) beamline 12-2 using a 20 μm X 40 μm beam at 0.9795 Å. Crystals were diluted and loaded onto a Mitegen MicroMesh 700/25 Loop. Frames were collected for 1.0 s with a 1.0° oscillation. The angular extent of diffraction and unit cell dimensions were assessed using ADXV.

### 2.4 Sample Preparation for Serial X-ray Experiments

Crystals were formed in batch, as described above, at a constant stir rate of 500 RPM. Further preparation was determined by delivery method. When using the MESH injector (Sierra *et al.*, 2012*a*), the microcrystal slurry was delivered as is, in a Hamilton syringe. When using the LCP injector (Weierstall *et al.*, 2014), crystals were mixed with viscogens, either PEO, LCP, or cellulose. For PEO mixtures, microcrystal slurry was combined with a viscogen consisting of 10% PEG + 10% PEO, and various ratios of crystal slurry to viscogen were tested. For LCP mixtures, crystal slurries were centrifuged, the supernatant was removed, with a minimal volume (100 μL) added back to suspend the crystals. The crystals were then mixed with Monoolein (9.9 MAG) in 1:1.5 mass-to-mass ratio using coupled glass syringes (Ishchenko *et al.*, 2016). For cellulose mixtures, crystal slurries were centrifuged, the supernatant was removed, and crystals were directly mixed with a 20% hydroxyethyl cellulose in a 1:9 crystal-to-cellulose ratio, as previously described (Sugahara *et al.*, 2017).

### 2.5 Serial X-ray Data Collection & Analysis

For the MESH & LCP XFEL datasets, we collected data at LCLS-MFX (Sierra *et al.*, 2019) on an MX170-HS Rayonix detector in 2-by-2 binning mode. Crystals were delivered to the XRD interaction point using either a MESH injector (Sierra *et al.*, 2012*a*, 2016) or an LCP injector (Weierstall *et al.*, 2014). Data were collected with a 3 μm beam at 9.5 keV energy, pulsed at 10 Hz, with a pulse duration of 40 fs on average. Powder diffraction patterns of silver(I) behenate were used to estimate the detector distance. The *cctbx.xfel* GUI was used for real-time feedback on the hit-rate and indexing-rate, as well as to submit processing jobs onsite. Data were indexed and integrated using *dials.stills_process*. Initial indexing results were used to refine the detector model, as well as crystal models (Brewster *et al.*, 2018). Refinement of the detector distance and panel geometry improved the agreement between measured and predicted spots. Data were then merged and post-refined using *cxi.merge*. Error estimates were treated according to the Ev11 method (Brewster *et al.*, 2018; Evans, 2011), wherein error estimates were increased using terms refined from the measurements until they could better explain the uncertainty observed in the merged reflection intensities. For the cellulose XFEL dataset, we collected data at SACLA (Ishikawa *et al.*, 2012) using a Diverse Application Platform for Hard X-ray Diffraction in SACLA (DAPHNIS) (Tono *et al.*, 2015) at BL2 (Tono *et al.*, 2019). Diffraction images were collected using a custom-built 4M pixel detector with multi-port CCD (mpCCD) sensors (Kameshima *et al.*, 2014). Data collection was supported by a real-time data processing pipeline (Nakane *et al.*, 2016) developed on Cheetah (Barty *et al.*, 2014). Identified hit images were processed in CrystFEL version 0.6.3 (White *et al.*, 2016). Diffraction spots were indexed by DirAx (Duisenberg, 1992). Intensities were merged by Monte Carlo integration with the *process_hkl* command in the CrystFEL suite with linear scale factors and per-image resolution cutoff. We note that the data collected at SACLA could not be processed using *dials.stills_process* due to spot shape irregularities that are an artifact of the mpCCD detector.

### 2.6 MicroED Sample Preparation

Samples for MicroED were prepared as previously described (Martynowycz *et al.*, 2019*a*). A 2 μL aliquot of crystals from the batch solution was applied to a glow-discharged Quantifoil Cu200 mesh R2/2 holey carbon grid. The grid was gently blotted from the back in an FEI Vitrobot for 10 s at 100% humidity and then vitrified in liquid ethane. Grids were stored in liquid nitrogen until further use. Prior to data collection, the grids were clipped and loaded into an FEI Versa FIB/SEM at liquid-nitrogen temperature and milled as previously described (Martynowycz *et al.*, 2019*a*). The grids were coated with a thin layer of amorphous platinum to increase the contrast during FIB/SEM imaging (Martynowycz *et al.*, 2019*b*). Large crystals (10-50 um) near the center of the grid square were identified using 2 kV SEM. Crystals were milled using a 30 kV gallium ion beam with a stepwise decreasing beam current as the sample slowly approached its final thickness of approximately 200 nm. The final 10 nm on either side of the crystalline lamellae were milled at 10 pA to polish the crystalline surface.

### 2.7 MicroED Data Collection & Analysis

Grids containing milled crystals were transferred into an FEI Arctica TEM operating at an accelerating voltage of 200 kV under liquid nitrogen. Crystalline lamellae were identified initially in an all-grid atlas taken at 155x magnification, where crystals were apparent as semitransparent areas suspended over a sharp, straight strip of empty area created by the milling process. Continuous rotation MicroED data were collected in diffraction mode over an angular wedge between −60° and 0° from the untilted orientation at a rotation rate of 0.3° / s. The camera length was set to 2055 mm, and frames were read out every 2 seconds. Data were recorded on a CetaD detector operating in rolling shutter mode with 2-by-2 binning. The camera length was calibrated using a molybdenum foil. MicroED data were converted from the FEI SER format to SMV for data analysis using in-house software that is freely available (https://cryoem.ucla.edu/). Data were indexed and integrated with XDS and scaled in XSCALE (Kabsch, 2010).

### 2.8 Model Refinement and Analysis

Data were reduced as described above. Initial phases were calculated by molecular replacement using Phaser, with PDB ID *4YUM* as the search model. Prior to initial atomic refinement, R-free-flags were carried over from PDB ID *4YUM* and random displacements (σ=0.5 Å) were applied to the atomic coordinates to help remove model bias. Next, iterative cycles of model building and further refinement were performed until the models reached convergence. Individual atomic coordinates, atomic displacement parameters (B-factors), and occupancies were refined using *phenix.refine* (Adams *et al.*, 2010; Afonine *et al.*, 2012). Automatic identification of ordered solvent was performed during the early cycles of model refinement. Models and maps were visualized and rebuilding steps were performed using *Coot* (Emsley *et al.*, 2010). Final structural models were visualized using PyMOL software, and were also used for ensemble refinement using *phenix.ensemble_refine* (Burnley *et al.*, 2012). Input parameters for ensemble refinement (*ptls*, *tx*, *wxray_coupled_tbath_offset*) were optimized for each dataset.

## 3. Results

### 3.1 Optimization of Batch Crystallization

Large CypA crystals (**Figure S1**), on the order of hundreds of microns, or even millimeters, are readily obtained by vapor diffusion methods (Fraser *et al.*, 2009; Keedy *et al.*, 2015); however, these crystals are too large for either microfluidic serial crystallography or MicroED. Therefore, we sought to optimize the production of microcrystals (on the order of 10-50 μm) in batch format, so that they could be easily delivered to the X-ray beam for serial crystallographic measurements using microfluidic sample injectors. As a first step towards this goal, we systematically explored the phase space of CypA crystallization in the vicinity of the conditions that yield large crystals (protein concentration in the range of 80-100 mg/mL with 20-25% (w/v) PEG-3350 as a precipitant and HEPES buffer at pH 7.5). We adapted the established CypA crystallization protocol (Fraser *et al.*, 2009) to a microbatch format (rather than vapor diffusion), and tested an array of conditions by varying protein and precipitant concentrations across the two axes of a 96-well crystallization plate. The lowest concentration of protein and precipitant led to the formation of large crystals that were ideal for data collection under traditional rotation conditions. In microbatch format, conditions that resulted in large, single crystals contained substantially lower protein and precipitant concentrations relative to vapor diffusion experiments that yield similarly sized crystals. Increasing protein concentration led to the formation of a greater number of smaller crystals; however, they tended to cluster together and displayed a needle-like morphology (**Figure 1**). High protein concentrations also led to a large variation in crystal size, which we sought to avoid since crystal monodispersity is desirable for serial crystallography experiments. In addition to modulating protein concentration, precipitant concentration was also varied. Increasing the concentration of the precipitant led to increased crystal density while maintaining better monodispersity. At the highest precipitant concentrations we tested, the protein tended to aggregate, rather than crystallizing. Given these characteristics, we found that we could consistently get dense crystal slurries when the final PEG-3350 concentration was near 20% (**Figure 1**). Increasing protein concentration beyond 35 mg/mL did not lead to appreciable increases in crystal density, so we chose a final protein concentration of 35 mg/mL and a final PEG-3350 concentration of 20% for further optimization.

**Figure 1.**
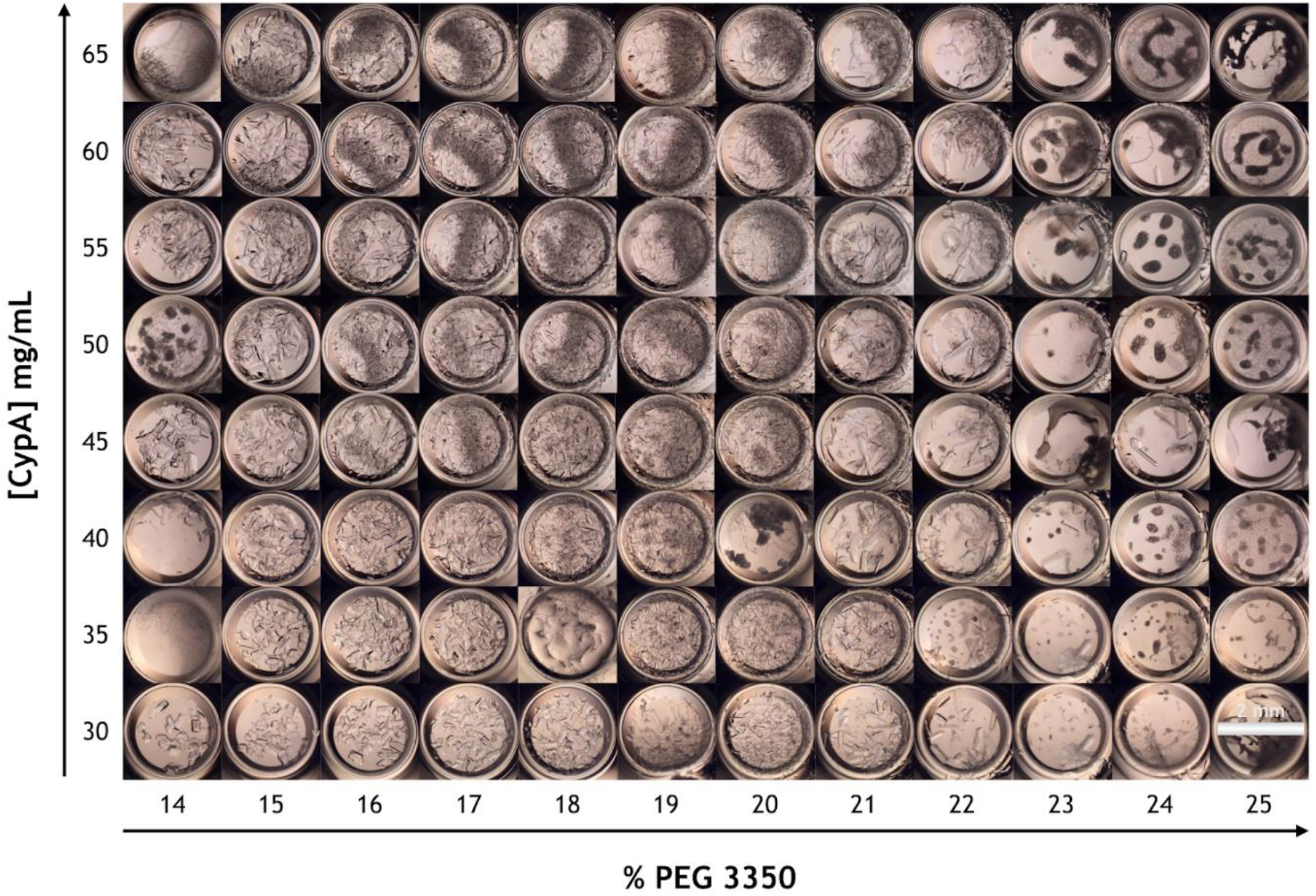
An array of images that illustrates the crystallization phase space of CypA. Concentrated solutions of CypA were mixed in a 1:1 volume ratio with solutions of varying concentrations of PEG-3350. Labels on the axes indicate final concentrations after mixing. CypA crystallizes readily in PEG-3350 solutions, however, the crystal size and morphology vary dramatically as a function of protein and PEG concentration. Specifically, at low [CypA] and low [PEG-3350] (bottom left corner), the crystals that form are few and large, while at high [PEG-3350] (right side), CypA aggregates and no crystals form. In the middle of the phase space, dense solutions of small crystals form.

After identifying ideal protein and precipitant concentrations using the microbatch method described above, our next goal was to scale up the microbatch procedure to produce crystal slurries at the milliliter scale. We developed a batch crystallization protocol in which 0.9 mL of CypA is stirred using a magnetic stir bar inside of a glass vial, and 0.6 mL of PEG-3350 solution (50% wt./vol.) is added dropwise to produce a solution with final protein and precipitant concentrations of 36 mg/mL and 20% (wt./vol.), respectively. Initially, we used a rotating mixer to mix the slurry by inversion, but we found that adding a stir bar yielded better results. Our batch stirring protocol also had the added benefit of improving monodispersity, decreasing crystal size, and increasing crystal density (crystals/mL) relative to the microbatch method. Increasing the final protein concentration above 36 mg/mL, and increasing the final PEG-3350 concentration above 20% did not further improve monodispersity, size, or crystal density; however, we discovered that modulating the stirring speed of the crystallization solution allowed us to control the formation of differently sized crystals (**Figure 2**). Within the range of stir-rates we tested (200-800rpm), we observed an increase in crystal density as stir rate increased, which was coupled to a decrease in the average crystal size. Crystals 50 μm or larger developed at lower stir rates, while crystals tended towards 10-20 μm at higher stir rates. In addition to modulating the density of the slurry, the stirring rate also affected monodispersity (Ibrahim *et al.*, 2015). At 200 RPM, we observed greater variation in the size of the crystals, while at higher stir rates the crystals were more monodisperse, but tended to clump together.

**Figure 2.**
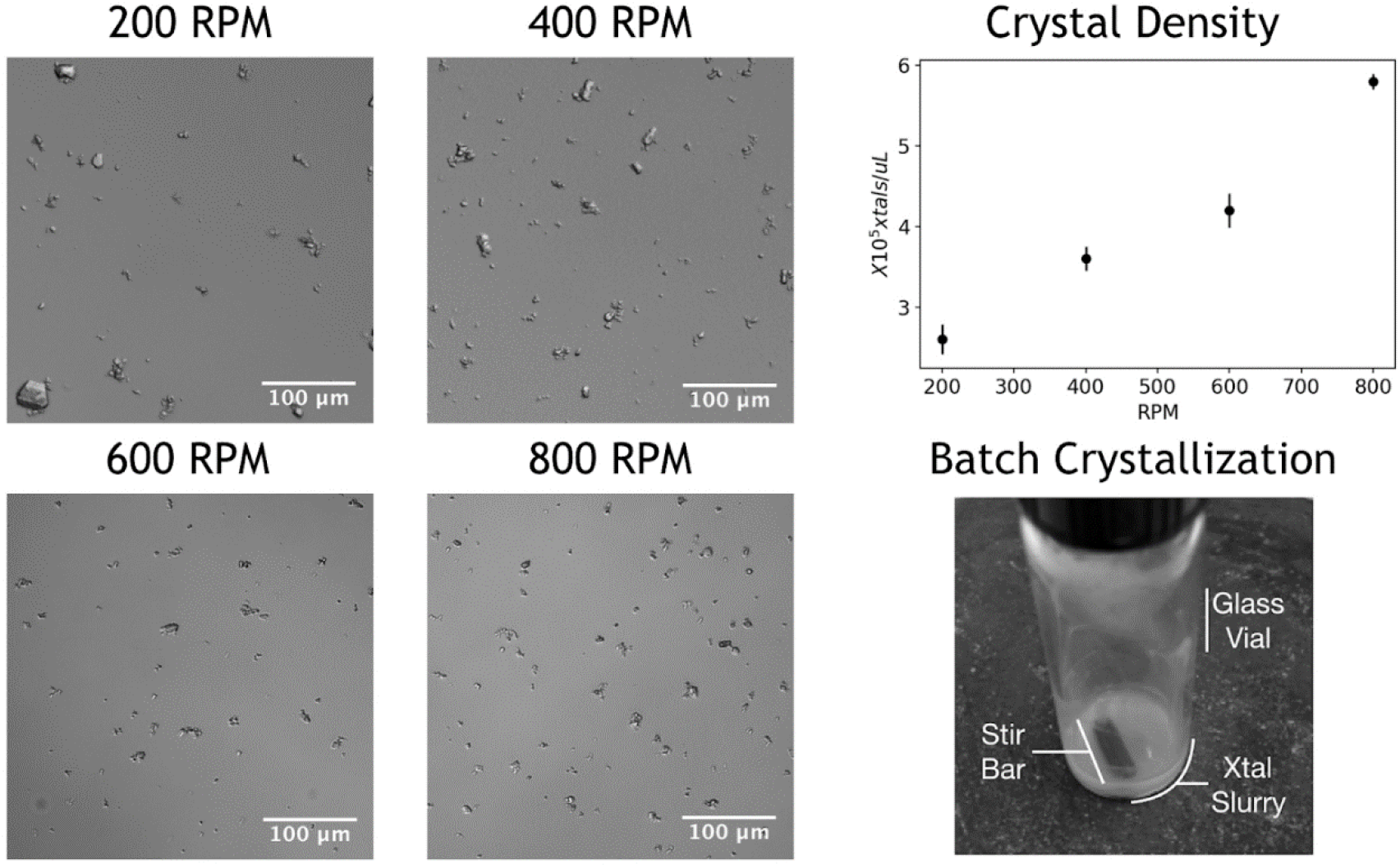
Images of microcrystals formed in batch with constant stirring. As stir rate increased, the average size of the crystals decreased, and the density of the slurry increased. This was confirmed by assessing crystal density using a hemocytometer.

Next, we tested how well the optimized microcrystals diffracted to ensure that the additional forces resulting from the stirring procedure did not degrade the quality of the crystal lattice. Batch-grown crystals were mounted on a polyimide mesh, and diffraction was measured at a synchrotron beamline (SSRL 12-2). These crystals were substantially smaller than crystals typically used for single-crystal X-ray crystallographic structure determination, however Bragg peaks were visible beyond 2.0 Å (**Figure S2**), and the indexed unit cell dimensions matched those of previously published CypA structures (Keedy *et al.*, 2015), indicating that the batch crystallization method does not have an adverse effect on the quality of the crystal lattice. Single crystals of this size (<50 μm) typically do not yield complete data sets at high resolution because of radiation damage considerations. It is becoming increasingly common to merge diffraction measurements from many small crystals, collected under traditional rotation conditions, but we did not pursue this mode of data collection for our batch-grown CypA microcrystals.

### 3.2 Serial X-ray Crystallography – Microfluidic Sample Delivery and Data Collection

We used CypA microcrystal slurries to perform several different types of serial crystallography experiments, all conducted in ambient atmosphere, in order to assess the performance of different methods of microfluidic sample delivery, and to determine whether the conditions created in the injector alters the outcome of the structural measurements. Specifically, we tested two different types of microfluidic sample injectors, one which utilizes the principle of electrospinning to form a microfluidic stream (Sierra *et al.*, 2012*a*), and another that performs high-pressure microextrusion of crystals embedded in a viscous material (Weierstall *et al.*, 2014). Because the microextrusion method requires that samples are extremely viscous, we tested three different viscogens as additives to our CypA samples. Thus, a total of four unique experimental conditions were explored.

The first injector system we implemented, in an experiment conducted at the MFX endstation of the XFEL at the Linac Coherent Light Source (LCLS), is referred to as the microfluidic electrokinetic sample holder (MESH). This device relies upon the principle of electrospinning to break the surface tension of the crystal slurry and drive it into a microjet as it exits the tip of a capillary (Sierra *et al.*, 2012*b*). Within the MESH system, gentle pressure from a syringe pump drives crystals through a capillary (250 μm i.d.) toward the X-ray interaction point, and application of approximately 3000 V of electrostatic potential (**Table 1**) across the sample stretches the liquid into a thin jet as it emerges from the capillary tip. Although our crystals were much smaller than 250 μm, they had a tendency to cluster together (**Figure 2**), so we used a capillary with a relatively large internal diameter to avoid clogging. A full description of the optimal operating parameters for our experiment using the MESH is provided in Table 1. During this experiment, the position and physical dimensions of the Taylor cone and microfluidic jet formed by electrospinning (**Figure S3**) had a tendency to fluctuate. However, by positioning the injector so the X-ray beam was at the approximate position where the liquid within the Taylor cone accelerated and became the jet, we were able to collect data at a hit rate of approximately 19% and an indexing rate of 63% (**Table 2**). Moving the injector so that the beam was pointed at the jet itself resulted in an unacceptably low hit rate (<1%), and moving the injector so that the beam was positioned at a more robust, but thicker, region of the Taylor cone resulted in extremely high background scattering due to the excessive volume of solvent in the beam path.

**Table 1.**
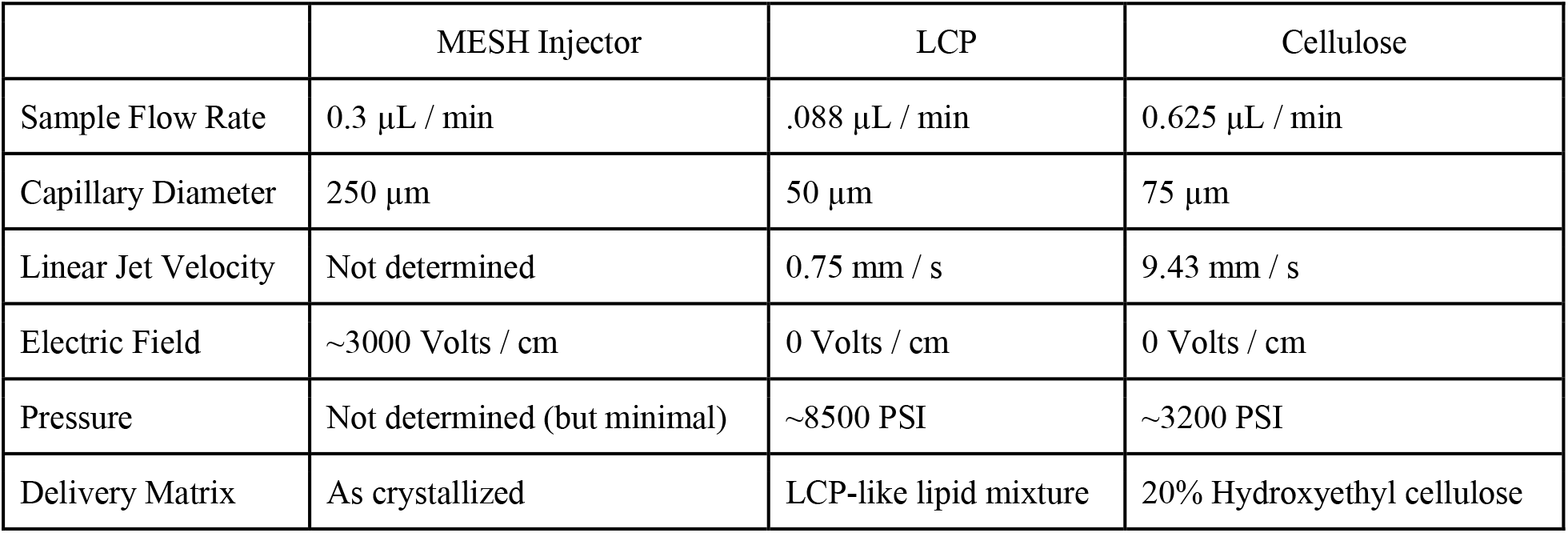
Sample injection parameters for serial XFEL experiments.

**Table 2.**
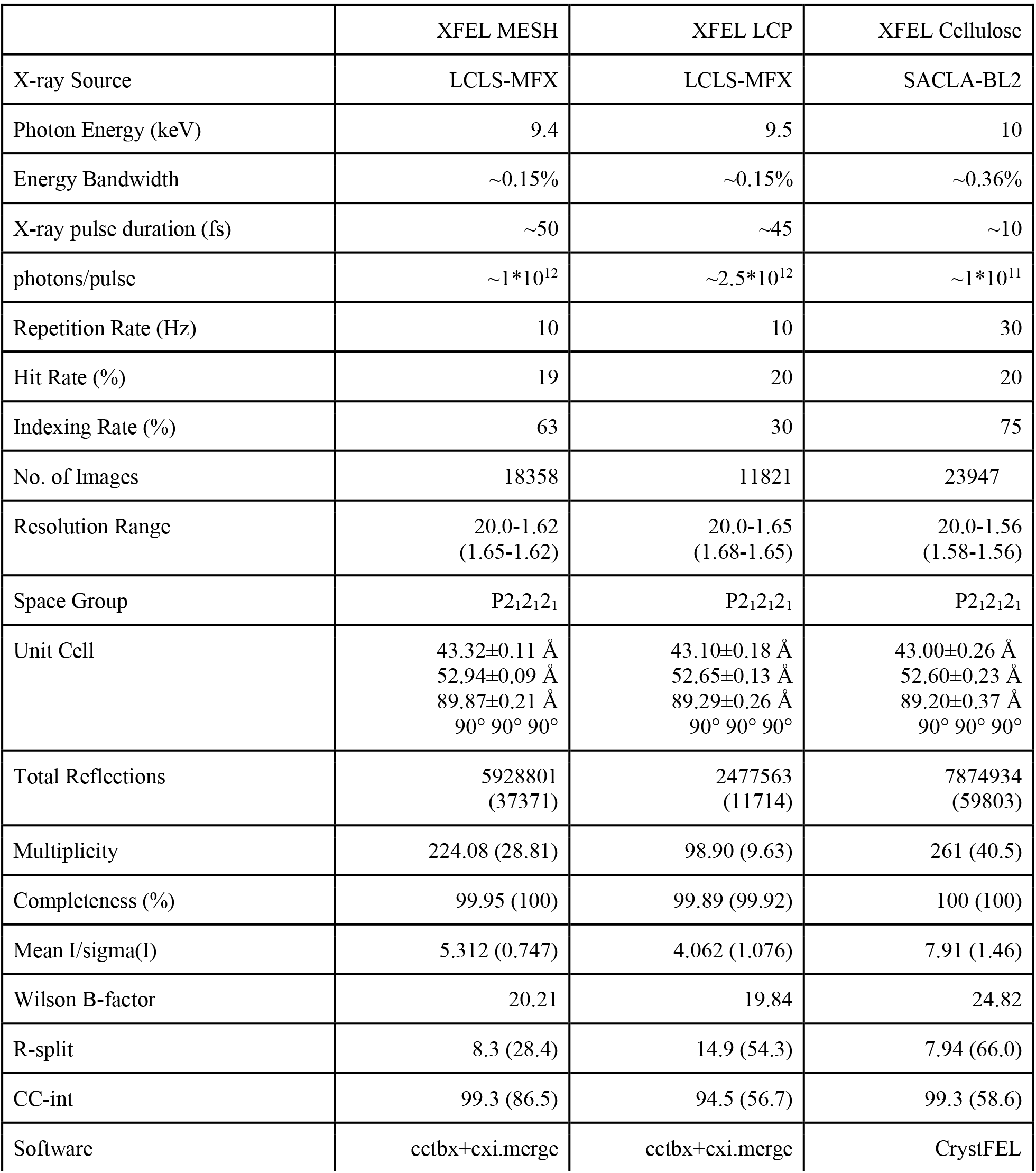
Crystallographic statistics for data collection. Statistics for the highest-resolution shell are shown in parentheses.

The second type of microfluidic sample injector we implemented for our experiments was a viscous extrusion system. Several variations on this device have been created, all of which operate using high pressure to extrude a viscous crystal slurry through a capillary, which is stabilized by a sheath gas to form a relatively slow-moving column of material as it exits the capillary at the X-ray interaction point (Weierstall *et al.*, 2014). Within the injector, crystals were exposed to pressures as high as 8500 PSI (**Table 1**), with no electrostatic potential. Given the high operating pressures of these injectors, we experienced no issues with clogging, so we used a 50 μm capillary to minimize sample consumption. Because these injector systems require that the sample be much more viscous than our CypA crystal slurries, their use required that we add viscogens, or “carrier media,” to our samples after crystallization but prior to injection. We experimented with three different types of carrier media: lipidic cubic phase (LCP) formed from monoolein, hydroxyethyl cellulose, and polyethylene oxide (PEO). Carrier media solutions were prepared independently and were mixed with crystal slurries to embed the microcrystals in the viscous material immediately prior to loading samples into the injector reservoirs. For LCP and PEO, the carrier media and crystal slurries we prepared in separate glass syringes, and mixed through a coupling device, while samples containing cellulose were prepared by mixing crystal slurry and carrier medium on a glass surface using a spatula *(Sugahara et al., 2015).* Visual inspections, using a microscope equipped with a cross-polarizer, confirmed that while it seems harsh, the process of mixing CypA microcrystals into viscous material does not visibly damage them. Crystal slurries prepared with LCP and PEO carrier media were delivered to the XFEL interaction point of the MFX endstation at the Linac Coherent Light Source (LCLS) using an injector device developed by (Weierstall *et al.*, 2014). Crystal slurries prepared with hydroxyethyl cellulose were delivered to the XFEL interaction point of the SPring-8 Angstrom Compact Linear Accelerator (SACLA), using an injector setup similar that used in studies Photosystem II (Suga *et al.*, 2017; Nango *et al.*, 2016; Kubo *et al.*, 2017). We observed that samples prepared with both LCP and hydroxyethyl cellulose formed microfluidic jets that were highly stable and maintained consistent physical dimensions for long periods of time, allowing efficient data collection. We obtained an average hit rate of 20% and an indexing rate of 30% for our experiment with LCP as the carrier medium, and we obtained an average hit rate of 20% and an indexing rate of 75% for our experiment with cellulose as the carrier medium (**Table 2**). On the other hand, and in contrast to reports by (Martin-Garcia *et al.*, 2017), we were unable to obtain useful data when PEO was used as the carrier medium. Samples prepared using PEO did not form stable jets, but instead formed droplets at the tip of the injector nozzle, which grew to a critical mass and then slowly dripped toward the X-ray interaction point and became unstable in the sheath gas. This problem persisted, despite considerable effort to optimize the sample and sheath gas flow parameters, and attempting to use both helium and nitrogen as the sheath gas.

In addition to assessing the data quality under different sample delivery conditions, we also wanted to determine whether the different types of sample delivery methods bias the orientation of the crystals as they are delivered to the X-ray beam (**Figure S4**). Because serial crystallography methods assume that crystals are delivered to the beam in random orientations in order to sample all of reciprocal space, the extent to which this is not true limits the efficiency of the experiment. We found that for the MESH data set, crystals do not appear to have an orientation bias as they are delivered to the X-ray beam, while the data set collected using LCP as a carrier medium for a viscous extrusion injector showed significant orientation bias.

### 3.3 Serial X-ray Crystallography – Data Quality and Atomic Structure are Robust Across Sample Delivery Strategies

Following data collection, the individual data sets were processed and the reduced data were compared, revealing that the high-quality diffraction typical of CypA crystals is robust across the different sample delivery methods that we implemented in our serial crystallography experiments. Raw diffraction images collected at LCLS (MESH and LCP conditions) were indexed and integrated using *dials.stills_process*, and the individual measurements were merged (with post-refinement) using *cxi.merge*. Raw diffraction images collected at SACLA (cellulose condition) were indexed, integrated, and merged using CrystFEL, following hitfinding with Cheetah. We used different software to process data obtained at different XFEL light sources because we observed that optimal software performance generally depends on idiosyncratic features of the experimental endstations, such as detector behavior and spectral characteristics of the X-ray pulses. The data sets were comprised of 18358, 11821, and 23947 indexed diffraction images each for the MESH, LCP, and cellulose samples respectively. The large number of indexed patterns used to construct each data set resulted in very high completeness and multiplicity, and CypA crystals diffracted to high resolution under all three delivery strategies (**Table 2**). The diffraction resolution of the data sets reported here fall within the range of resolutions reported for CypA structures solved using large crystals and rotation geometry, and the modest differences in maximum resolution (1.65-1.56 Å) between the data sets are likely due to differences in the number of indexed patterns contributing to each data set, rather than to significant differences in the quality of diffraction under the three distinct sample delivery conditions. Statistical metrics reflecting the measurement precision and strength of the diffraction signal were favorable for all data sets, indicating that none of the delivery methods compromised the integrity of the crystal lattice (**Table 2**).

Using the reduced data sets, we performed molecular replacement to calculate initial phases, followed by iterative cycles of model building and atomic refinement to determine the structure of CypA under each of the different sample delivery conditions. After molecular replacement and before the initial cycle of manual model building, we applied random perturbations (σ = 0.5 Å) to the atomic coordinates and refined them against the X-ray data in order to eliminate any effect of model bias that might arise from using the same molecular replacement search model for the three independent structures. We performed iterative rounds of model building and atomic refinement until the procedure reached convergence, and found that the models we obtained from each of the three experiments were of comparable statistical quality in terms of their fit to the experimental data and overall geometry (**Table 3**). We discovered that the method of sample delivery in each of the three serial crystallography experiments has minimal impact on the average structure of CypA; however, the three individual structures are not identical.

**Table 3.** Statistics for X-ray model refinement. Statistics for the highest-resolution shell are shown in parentheses.

|  | XFEL MESH | XFEL LCP | XFEL Cellulose |
| --- | --- | --- | --- |
| Resolution Range | 19.91-1.62<br>(1.72-1.62) | 17.43-1.65<br>(1.75-1.65) | 19.9-1.56<br>(1.64-1.56) |
| Unique Reflections | 26445 (4324) | 25034 (4076) | 29528 (4159) |
| Reflections used in Refinement | 25,613 (4189) | 24247 (3950) | 28,598 (4027) |
| Reflections used for R-free | 832 (135) | 787 (126) | 929 (132) |
| R-work | 0.1362 (0.2398) | 0.1434 (0.2316) | 0.1348 (0.2376) |
| R-free | 0.1569 (0.2455) | 0.1671 (0.2367) | 0.1509 (0.2804) |
| Number of non-hydrogen atoms | 1522 | 1558 | 1551 |
| macromolecules | 1383 | 1399 | 1401 |
| Protein residues | 163 | 163 | 163 |
| RMS (bonds) | 0.005 | 0.004 | 0.015 |
| RMS (angles) | 0.844 | 0.656 | 1.298 |
| Ramachandran favored (%) | 96.89 | 96.89 | 96.89 |
| Ramachandran allowed (%) | 3.11 | 3.11 | 3.11 |
| Ramachandran outliers (%) | 0.00 | 0.00 | 0.00 |
| Rotamer outliers (%) | 0.68 | 2.68 | 0.00 |
| Clashscore | 3.24 | 1.43 | 1.78 |
| Average B-factor | 26.29 | 26.78 | 29.62 |
| macromolecules | 24.78 | 25.29 | 28.02 |
| PDB ID | 6U5C | 6U5D | 6U5E |
| Ensemble Refinement |  |  |  |
| R-work | 0.1241 | 0.1304 | 0.1296 |
| R-free | 0.1477 | 0.1517 | 0.1512 |

Pairwise alignment of the three structures and comparison of atomic coordinates revealed that for each pair of structures, the root-mean-squared deviation (RMSD) of atomic positions is less than 0.1 Å (**Table 4**). Conformational heterogeneity of a key network of residues in CypA that extends from the core of the protein to the active site has been studied previously using ambient-temperature crystallographic experiments. Rotameric exchange of residues in this network, which includes R55 (the catalytic residue), M61, S99, and F113, is required for enzymatic turnover (Eisenmesser *et al.*, 2005; Fraser *et al.*, 2009). Notably, all three of our serial datasets revealed evidence for multiple conformations of these residues (**Figure 3**). Small differences between the structures existed as differences in rotamers (or mixtures of rotamers) for side chains with generally weak electron density, such as Met61.

**Table 4.** All-atom RMSD values for comparison of the three serial crystallography structures.

| Pair | RMSD |
| --- | --- |
| MESH/LCP | 0.048 Å |
| MESH/Cellulose | 0.063 Å |
| LCP/Cellulose | 0.069 Å |

**Figure 3.**
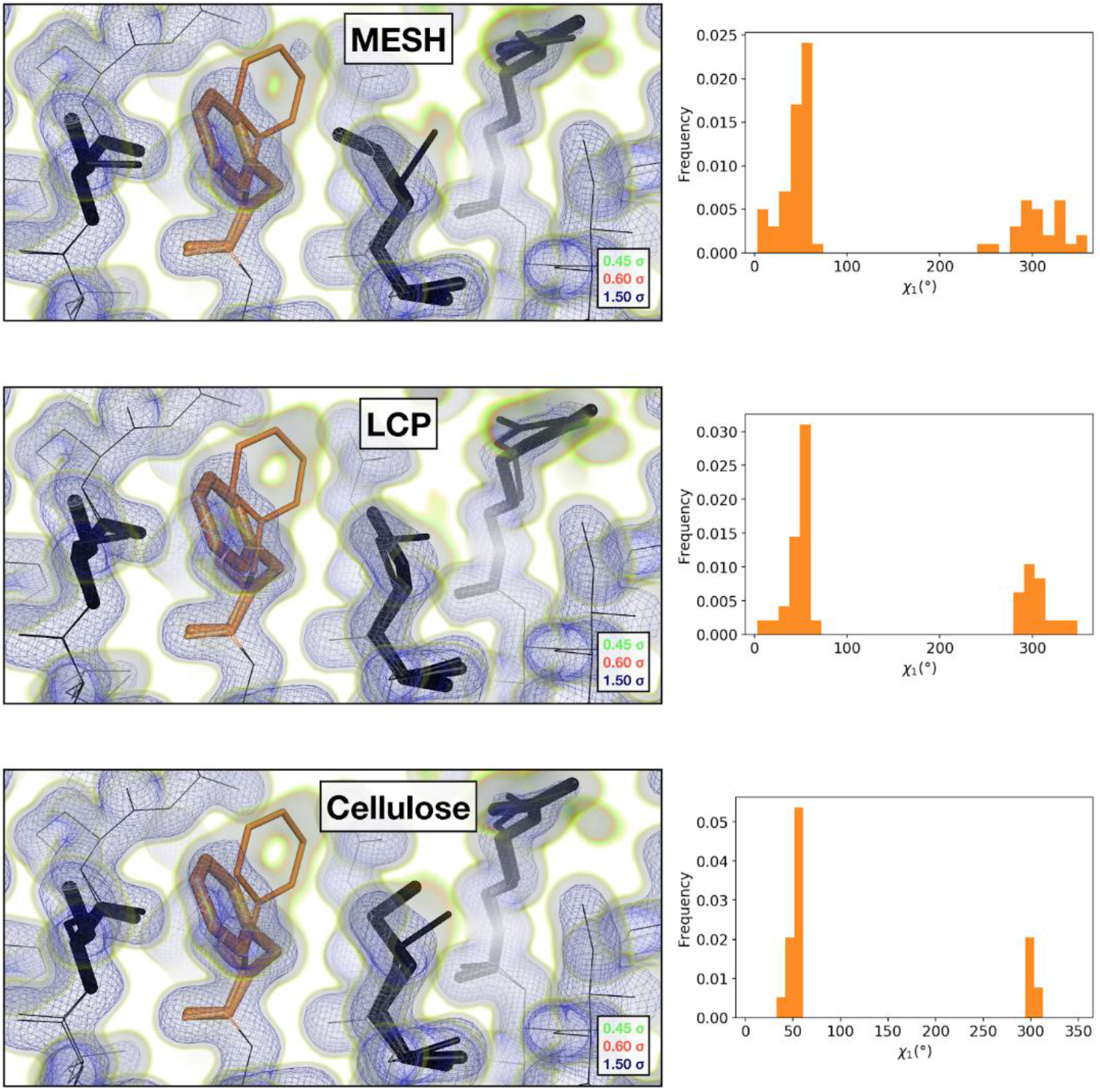
Comparison of the 2mFoFc map and the refined multi-conformer model produced from each serial XFEL experiment. Maps were visualized at multiple contour levels to show evidence for alternative conformations. Following multi-conformer refinement, ensembles were generated from each model using *phenix.ensemble_refine*. In the right panel, a histogram of the Chi1 angles for residue 113 is plotted for the ensemble. Multi-conformer models plus maps, and the distribution of Chi1 angles across the ensemble models are similar for all three XFEL data sets.

Furthermore, CypA’s structure did not appear to be significantly perturbed by either the electric field within the MESH jet, or by the high pressures within the LCP injector used for the LCP and Cellulose datasets (**Figure 3**). Local features within the models matched the maps well, with only subtle differences noticeable in the maps. Model statistics were similar across all three datasets, only the average B-factor differed between appreciably between datasets (**Table 3**). Comparing the normalized atomic B-factors of atoms within each structure (**Figure S5**) revealed that the increase in the average was not due to any localized change in conformational heterogeneity, but instead resulted from a global increase in the refined B-factors. These global differences in atomic B-factors across structures could be due to varied perturbation of the crystal lattice (but not the molecular structure) that results from exposure to different sample delivery conditions, or due to small differences in the data processing parameters. When the structure was expanded to an ensemble, the resulting multiconformer models settled into nearly equivalent minima, confirming the similarity of the three datasets (**Figures S5 & S6)**.

### 3.4 MicroED – Grid Preparation and Data Collection

In order to obtain a CypA microcrystals on copper grids, suitable for MicroED data collection, we tested several sample preparation strategies. The ideal crystal thickness for MicroED samples is approximately 300-500 nm (Martynowycz *et al.*, 2019*b*), which is substantially smaller than any crystal that is visible using light microscopy. First, we prepared grids using a CypA microcrystal slurry containing visible crystals on the order of 10 um, with the hope that this sample would also contain much smaller crystal fragments that would be acceptable for data collection. We examined this sample in the microscope and observed only large (several microns or larger) microcrystals on the grid. As a next step, we attempted to reduce the size of the CypA microcrystals using several physical agitation methods, including vortexing and crushing the crystals using either a pipette tip or glass beads. Samples exposed to physical agitation were used for grid preparation and were examined under the microscope, again revealing an absence of suitably sized crystals for data collection. Attempts to improve the grid preparation by changing the glow discharge and blotting methods also did not result in suitable samples. We hypothesize that difficulties in preparing grids with sub-micron sized CypA crystals result from a combination of the crystals’ surface properties and the strong lateral forces that are introduced by the blotting process, which could pull small crystals off of the grid. We note that we used only grids with amorphous holey carbon supports, and did not attempt to prepare samples using grids with more exotic support materials, such as gold or graphene oxide.

Because we were unsuccessful in preparing samples for MicroED using traditional methods of applying crystals to grids, we turned to a method that utilizes a focused ion beam (FIB) to mill larger crystals down to an appropriate thickness. We observed that crystals larger than several microns were able to stick to the holey carbon grids, we prepared a frozen grid with crystals that were approximately 5-10 um. Prior to MicroED data collection in the transmission electron microscope (TEM), the frozen samples were loaded into a scanning electron microscope (SEM) equipped with a gallium ion FIB. A single crystal, approximately 20-30 μm in all dimensions was identified in the SEM and was subsequently FIB-milled to form a crystalline lamella that was approximately 200 nm thick (**Figure 4**). The grid containing the lamella was then transferred to the TEM for MicroED data collection. Three separate data sets were collected from unique regions of the lamella, two of which were merged to produce the final reduced data set that was used for structure determination. Inclusion of the third data set degraded the quality of the merged data. Also, because the rotation range of the microscope stage is restricted, and the crystal orientation was the same for all three data sets due to the fact that they were collected from a single lamella, the final merged data set suffered from a missing wedge of reciprocal space (**Figure 4 and Supporting Figure 8**) and had an overall completeness of 86% at 2.5 Å resolution. Additional information about the quality of the merged MicroED data is provided in Table 5.

**Table 5.**
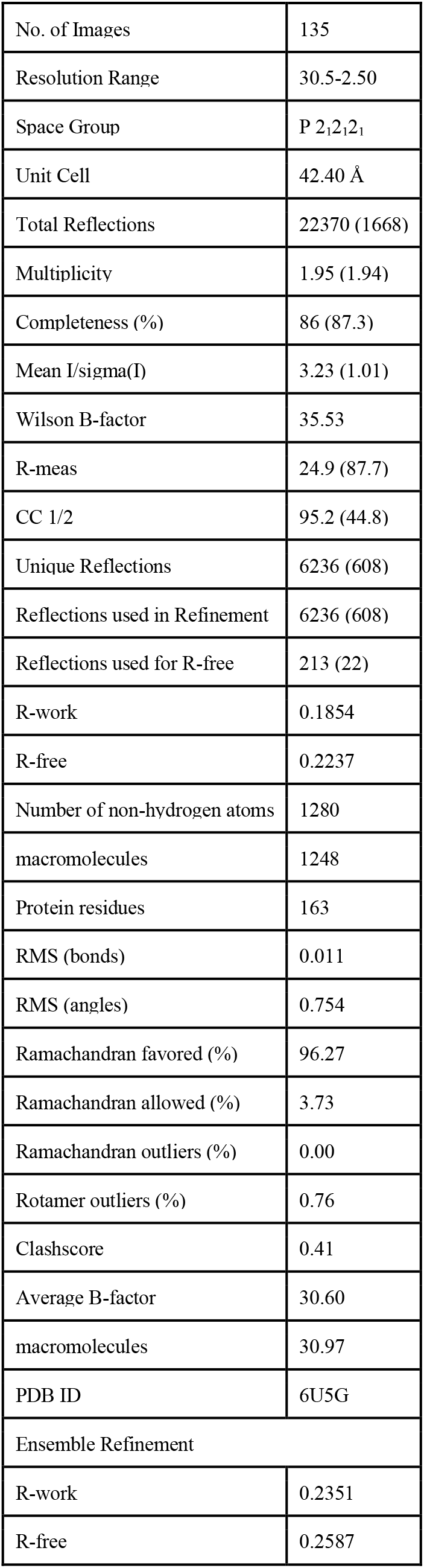
Crystallographic statistics for MicroED data. Statistics for highest-resolution shell are shown in parentheses.

|  |  |
| --- | --- |
| No. of Images | 135 |
| Resolution Range | 30.5-2.50 |
| Space Group | P 2<sub>1</sub>2<sub>1</sub>2<sub>1</sub> |
| Unit Cell | 42.40 Å |
| Total Reflections | 22370 (1668) |
| Multiplicity | 1.95 (1.94) |
| Completeness (%) | 86 (87.3) |
| Mean I/sigma(I) | 3.23 (1.01) |
| Wilson B-factor | 35.53 |
| R-meas | 24.9 (87.7) |
| CC 1/2 | 95.2 (44.8) |
| Unique Reflections | 6236 (608) |
| Reflections used in Refinement | 6236 (608) |
| Reflections used for R-free | 213 (22) |
| R-work | 0.1854 |
| R-free | 0.2237 |
| Number of non-hydrogen atoms | 1280 |
| macromolecules | 1248 |
| Protein residues | 163 |
| RMS (bonds) | 0.011 |
| RMS (angles) | 0.754 |
| Ramachandran favored (%) | 96.27 |
| Ramachandran allowed (%) | 3.73 |
| Ramachandran outliers (%) | 0.00 |
| Rotamer outliers (%) | 0.76 |
| Clashscore | 0.41 |
| Average B-factor | 30.60 |
| macromolecules | 30.97 |
| PDB ID | 6U5G |
| Ensemble Refinement |  |
| R-work | 0.2351 |
| R-free | 0.2587 |

**Figure 4.**
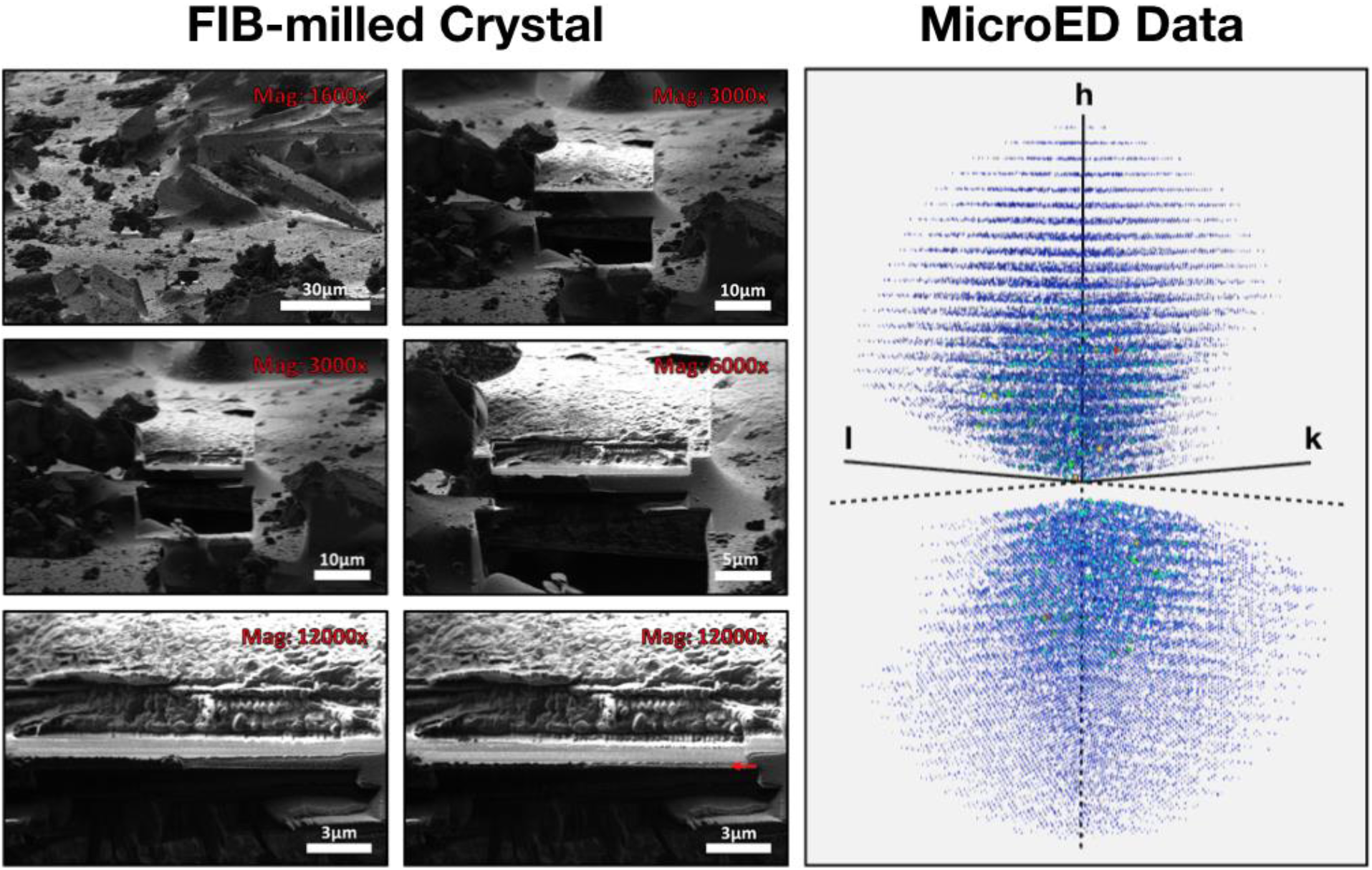
MicroED Data collection. CypA crystals were deposited on a copper grid with an amorphous carbon support material and frozen in vitreous ice (left). A single crystal is shown at various stages of the FIB milling process, progressing from the upper left to the lower right. The final crystalline lamella is denoted with a red arrow in the bottom right image. Also shown is the intensity weighted reciprocal lattice (right) representing the MicroED data that was collected from the single crystal shown in the left panel.

### 3.5 CypA Structure Determination from MicroED Data

After merging the integrated MicroED data to obtain a high-quality data set, we implemented the exact same structure determination procedure as was used for analysis of the serial X-ray diffraction data sets. Specifically, we performed molecular replacement followed by application of random coordinate perturbations, and then iterative model building and refinement until the Rwork and Rfree values converged and no additional improvements to the model could be made. Analysis of the CypA crystal structure determined by MicroED revealed two notable features.

First, during the indexing stage of the data reduction procedure, we observed that the unit cell had unusual dimensions (**Figure 5**). Specifically, while the crystallographic a and c axes match well to those of other CypA structures determined at cryogenic temperatures, the b-axis is approximately 1% longer than the corresponding axes in typical CypA structures determined at “physiological” (>260 K) temperatures. Because of indexing challenges resulting from the slight incompleteness of the data, such as the inability to observe reciprocal lattice points along the principal k and l axes within the 60° wedge that we measured (**Figure 4**), we took several additional steps to ensure that the unit cell was indexed accurately and that the space group symmetry was correctly assigned. To address the possibility that an optical distortion in the microscope or challenges related to the flatness of the Ewald sphere (Clabbers & Abrahams, 2018) could lead to incorrect measurement of unit cell dimensions, we performed a structure refinement procedure that also simultaneously refined the coordinates and the lengths of the unit cell axes. This “unit cell refinement” procedure resulted in refined unit cell dimensions that were essentially the same as the input, indicating that the indexing converged to the correct solution. To confirm that the elongation of the b-axis does not also break the crystallographic P2_1_2_1_2_1_ space group symmetry, we reduced the raw data three separate times in space group P2_1_. In each of these three data sets, the two-fold symmetry operation was preserved along a different crystallographic axis (i.e. P2_1_11, P12_1_1, and P112_1_ relative to the parent P2_1_2_1_2_1_). Refinement of the CypA structure against the data with lower symmetry produced models with worse overall quality than when the data were reduced in space group P2_1_2_1_2_1_, confirming the validity of the space group assignment.

**Figure 5.**
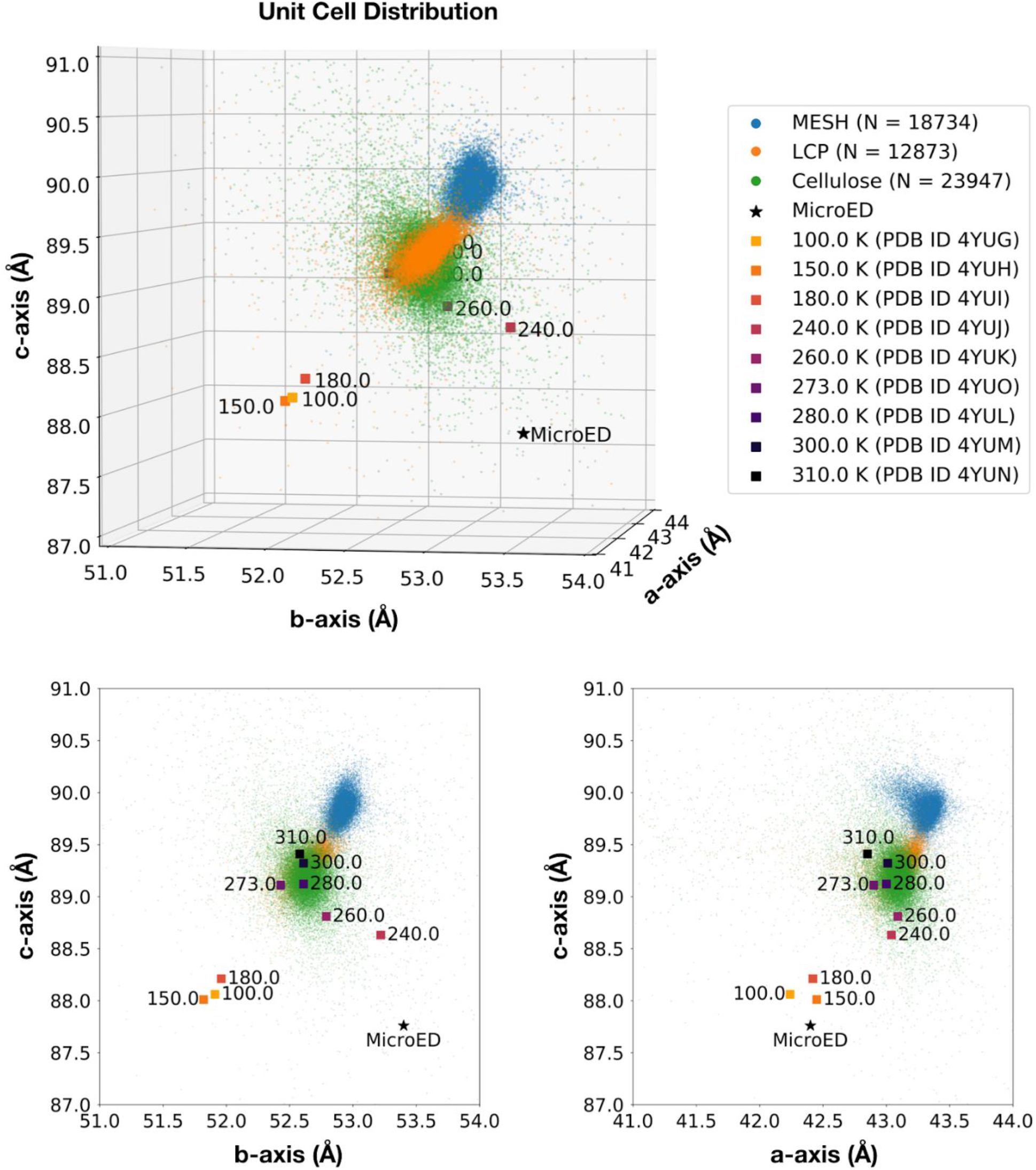
Comparison of unit cell dimensions across data collection strategies. Published structures are provided as a reference for the effect of temperature upon unit cell dimensions. The unit cells measured using serial XFEL experiments resemble data from published room temperature structures. A fib-milled crystal used for MicroED revealed dimensions that were unique from the unit cell compression normally seen in cryogenic X-ray data.

The second notable observation that we made about the crystal structure determined by MicroED is that, while the unit cell is distorted relative to other CypA structures, the structure of the molecule within the unit cell is essentially the same as for X-ray structures that were determined using cryocooled crystals (PBD ID *3K0M*), RMSD = 0.22 Å. Structures of CypA determined from cryocooled crystals, using both X-ray and MicroED, lack key conformations that are visible in their ambient temperature counterparts. In particular, ambient temperature structures of CypA reveal alternative conformations of an important network of amino acid side chains (the catalytic residue R55, as well as M61, S99, and F113), while structures determined using cryocooled samples, including the MicroED structure presented here, reveal only a single conformation of these side chains (**Figure 6**). These alternative conformations are visible in electron density maps derived from the X-ray data, even when the resolution is reduced to match that of the MicroED data (**Supporting Figure 9**).

**Figure 6.**
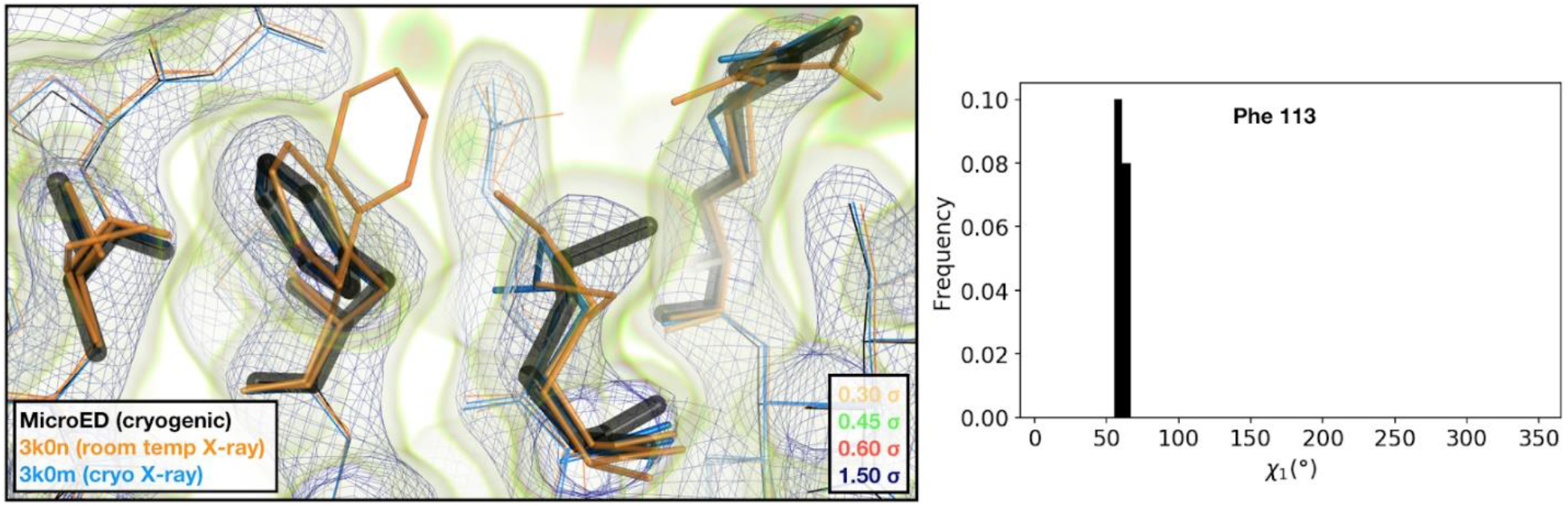
Visualization of the 2mFoFc map and the refined model measured from a fib-milled crystal using MicroED. The conformation of residues coupled to the catalytic site resembles structures previously solved under cryogenic conditions using X-ray crystallography (PDB ID *3K0M*). For some regions of the structure, the cryogenic X-ray and MicroED structures are indistinguishable. A previously published multi-conformer model produced from data acquired at room temperature is provided for comparison (PDB ID *3K0N*). Following refinement, ensembles were generated using *phenix.ensemble_refine*. In the right panel, a histogram of the Chi1 angles for residue 113 is plotted for the ensemble. All members of the ensemble adopted the same rotameric position as previous cryogenic structures.

### 3.6 Effect of Experimental Conditions on Unit Cell Dimensions

Comparing the structures that we determined using different microcrystallography techniques revealed that the unit cell dimensions of the CypA crystals were noticeably affected by the conditions required for each of the different experiments (**Figure 5**). The three serial X-ray datasets closely resemble previous room-temperature data collected under traditional rotation conditions. Additionally, we observed that for sample delivery methods that involve embedding the crystals in a viscous carrier medium (LCP or cellulose), the unit cell tended to be slightly smaller than for experiments that do not require such additives (MESH). Average unit cell dimensions for the LCP and cellulose data sets were approximately 0.5-0.7% smaller than for the MESH data set (roughly twice the value of the standard deviations of the corresponding unit cell distributions). We hypothesize that the observed shrinkage of the unit cell could be due to crystal dehydration. The MicroED data revealed a highly unusual unit cell, with an expanded b-axis relative to any other CypA structure that has been reported (**Figure 5**). The unit cell was different from both ambient temperature (PDB ID *3K0N*) and cryogenic (PDB ID *3K0M*) X-ray structures, and matches most closely to a structure determined at 240 K (PDB ID *4YUJ*). We hypothesize that the unusual unit cell observed in the MicroED experiment could be the result of cooling the crystals in ethane, rather than nitrogen, or could be caused by the grid blotting procedure or FIB-milling, both of which are unique to MicroED. Despite the small variations in crystal packing that cause changes in unit cell parameters for structures determined using different methods, the refined coordinates of CypA molecules themselves are quite consistent.

## 4. Discussion

The ability to measure diffraction signals from ever smaller crystal samples has enabled a variety of new and innovative experiments in macromolecular crystallography, however there is still a relative absence in the literature of practical guidelines for optimizing microcrystallography experiments. The work we present here attempts to address this knowledge gap, by providing a detailed description of how we optimized the growth of cyclophilin A (CypA) microcrystals and measured their diffraction using two emerging microcrystallography techniques, serial XFEL crystallography and microcrystal electron diffraction (MicroED). Our results compare and contrast serial X-ray and MicroED methodologies, and highlight some important considerations and pitfalls that might be encountered during preparation of microcrystalline samples for the respective experiments. This case study provides a roadmap for experimenters who are interested in performing structural measurements using crystalline samples with dimensions on the scale of nanometers to microns.

For decades, macromolecular crystallographers have strived to grow large (hundreds of microns) single crystals that can be used for crystallographic measurements using rotation X-ray methods, but new data collection methods such as serial X-ray crystallography and MicroED require the reliable formation of crystals that are much smaller, typically hundreds of nanometers to tens of microns. Precise control of crystal size over this range is challenging, and others have developed methods that employ specialized equipment for *in situ* light scattering measurements to evaluate crystal size in real time (Baitan *et al.*, 2018; Schubert *et al.*, 2017) and halt crystallization as it progresses. Instead, our work with CypA demonstrates a simple, alternative method for controlling the size of crystals during batch growth. Starting from crystallization conditions identified by microbatch screening in 96-well plates, we scaled up the crystallization volume and introduced agitation (by stirring) to control the crystal size. We observed that at higher stir rates (i.e. more agitation) crystals tend to be smaller and more concentrated in the resulting slurry. We speculate that stirring fractures the crystals when they reach a critical size, which exerts control over the crystal dimensions and also actively introduces seeds into the slurry. For CypA, we found that batch crystallization with stirring could be used to generate relatively monodisperse crystal slurries, in milliliter volumes, with crystal sizes in the range of microns to hundreds of microns. We believe that the batch stirring protocol is likely to be useful for a variety of crystal systems beyond CypA, however it may have limited utility for crystal systems that are more susceptible to physical damage.

With the ability to create large batches of CypA crystals, we could perform serial X-ray crystallography experiments at XFEL lightsources, which generally consume a large amount of sample. We utilized CypA microcrystal slurries, prepared in an identical fashion, to evaluate several commonly used, injector-based sample delivery strategies, including both electrospinning and viscous extrusion using two types of crystal carrier media. These delivery strategies exposed the crystals to extreme experimental conditions, including strong electric fields, high pressures, and unusual carrier media. Crystal structures determined using each method revealed how conditions imposed by the different sample delivery systems perturbed either the crystal lattice or the protein structure. We observed that the different sample delivery methods do produce measurable differences in the distributions of unit cell axis lengths for CypA microcrystals. Crystals measured using the MESH device (electrospinning) tended to have longer a, b, and c axes than crystals measured using viscous carrier media, which might result from dehydration of the crystals by the viscogens (LCP or cellulose), which reduce the relative humidity of the crystallization mother liquor. Additionally, we observed that crystals delivered using the MESH (electrospinning) device tended to be oriented more randomly than crystals delivered in a viscous carrier such as LCP (**Figure S4**). In the case of our CypA crystals, the dihedral space group symmetry prevents the crystals from having a dipole moment that could cause them to assume a preferred orientation in the electric field introduced by the MESH injector. On the other hand, the slightly elongated crystal morphology led to orientation bias, likely due to shear forces resulting from the flow of the highly viscous liquid. We expect that crystals with different properties, such as polar space group symmetry or more isotropic morphologies, would have different behaviors with respect to preferred crystal orientation in the various injector systems. Despite differences in unit cell parameters and preferred crystal orientations, the overall quality of the reduced data sets resulting from each of the serial X-ray experiments were generally equivalent (**Table 3**), as were the resulting atomic models (**Table 4**). We also utilized a multi-conformer ensemble refinement approach as a way to assess the level of heterogeneity (model variance) that was present in each of the data sets. Our analysis focused on a network of catalytically-important residues, which are known to be dynamic (Eisenmesser *et al.*, 2005; Fraser *et al.*, 2009). We observed that the refined ensembles reflect a similar level of heterogeneity across the different structures (**Figure S6**), which is generally supported by the correlations in the B-factors derived from standard refinements (**Figure S5**). We did, however, observe that the ensemble derived from the MESH data shows enhanced heterogeneity relative to the other two data sets for a loop region including residues 69-74 (**Figure S7**). The conformation of this loop is stabilized by a key charged residue (R69) (Caines *et al.*, 2012), which may be perturbed by the electric field. While it has been shown that electric fields can be used to perturb conformational dynamics in proteins (Hekstra *et al.*, 2016), we expect that the effect should be minimal in our MESH experiment, because the crystals are randomly oriented relative to the applied electric field, and the field is more than two orders of magnitude less than those that are intentionally used for perturbing conformational dynamics (Hekstra *et al.*, 2016). Our results show that the choice of microfluidic sample delivery method has a minimal effect on the static crystal structure of CypA. Consequently, the choice of sample delivery method for a serial X-ray crystallography experiment should be selected based on practical considerations related to the experiment, such as the requirement for a laser perturbation or mixing in a time-resolved experiment.

In addition to serial X-ray crystallography, MicroED also offers the ability to determine macromolecular structures at high resolution using very small crystals with moderate solvent content. In stark contrast to serial X-ray crystallography experiments, which require hundreds of milligrams of protein and milliliters of microcrystal slurries, MicroED lies at the other extreme, allowing crystal structure determination with as little as a single microcrystal. Additionally, MicroED experiments have a significant advantage in that they are much more accessible than experiments performed at XFELs, and require a substantially lower investment of time and resources. Using CypA crystals derived from our batch protocol, we encountered several challenges in preparing appropriately-sized microcrystal samples on grids for measurement in the TEM. MicroED requires extremely small crystals, ideally less than 500 nm thick (Martynowycz *et al.*, 2019*b*), and we encountered difficulties in getting such small CypA crystals to remain on the grids after blotting away excess solvent. This may be due to the specific surface chemistry of CypA crystals or may be a more general trend of high solvent content crystals. As a result, we turned to a recent development in sample preparation that is widening the scope of MicroED by enabling measurements from crystals that are initially tens of microns thick, by utilizing a FIB-milling process to machine large crystals into thin lamellae that are optimal for MicroED measurements (Martynowycz *et al.*, 2019*a*). The FIB-milling procedure allowed us to determine the structure of CypA by MicroED using a single crystal that was initially (before milling) similar in size to those which we used for serial X-ray experiments. We observed that the MicroED crystal structure of CypA had a slightly distorted unit cell relative to other reported CypA structures, and we hypothesize that this could be due to the damage from either blotting or FIB-milling, however more rigorous studies will be required to evaluate the specific effects of these sample prep procedures on MicroED structures. The sensitivity of the CypA crystals during preparation for MicroED could be related to their high solvent content. Our structure of CypA has the highest solvent content of any non-membrane protein MicroED crystal structure determined to date using three-dimensional crystals (**Figure 7**), demonstrating how improved sample preparation is expanding the technique to include more challenging crystal systems.

**Figure 7.**
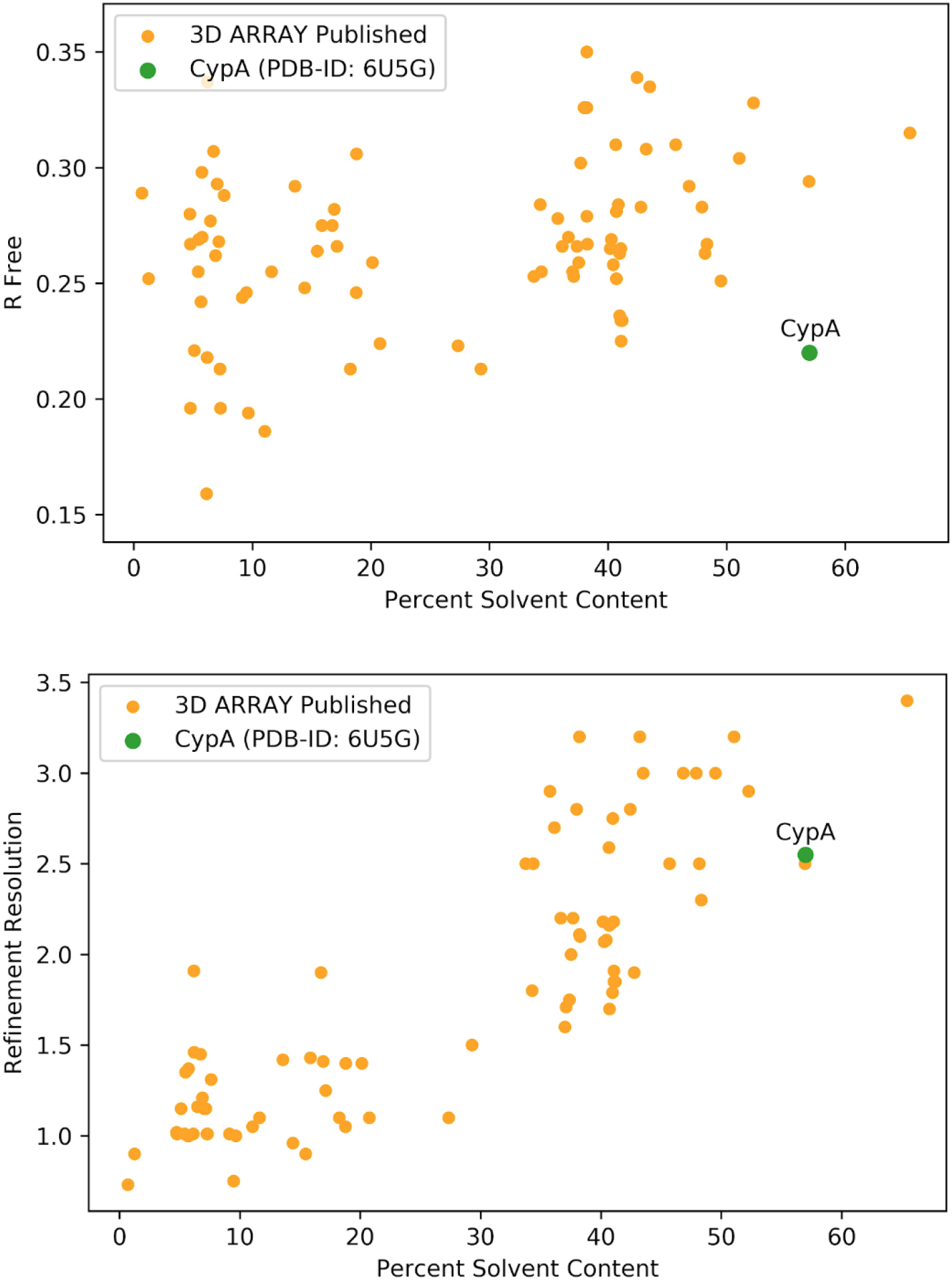
Survey of MicroED datasets deposited on the PDB. Structures determined from 3D crystals are shown as orange points, and CypA is shown as a green point. The highest solvent content point is 3J7T, which is a membrane protein.

Our evaluation of modern protein microcrystallography techniques reveals that MicroED and serial X-ray crystallography are complementary methods for structural biology (Zatsepin *et al.*, 2019). The optimal experimental method for a microcrystallography experiment will depend upon various aspects of the macromolecular system that is being studied. For determination of static, low-energy macromolecular structures, MicroED has substantial advantages over serial X-ray crystallography in terms of sample prep requirements, quantity of material required, and ease of data collection. However, while serial X-ray crystallography experiments require large amounts of sample, specialized equipment that is only available at select X-ray lightsources, and substantial optimization of sample delivery parameters, they also have their advantages. Importantly, serial X-ray measurements are performed at ambient temperature and can reveal physiological conformational ensembles of the crystallized molecules. On the other hand, we compared our MicroED structure to a cryogenic X-ray structure of CypA (*3K0M*), and observed that it was nearly identical and suffered from the same temperature-dependent reduction in conformational heterogeneity (**Figure 6**). Because cooling rate is related to crystal size (Halle, 2004), it remains to be seen whether MicroED experiments using very small crystals (hundreds of nanometers) might capture a more physiological conformational ensemble. Our data do not shed light on this question, since the crystals used in our experiments were approximately 20-30 μm at the time of freezing, before they were FIB-milled to an appropriate thickness. Finally, we note that for CypA, as well as other examples from the literature including lysozyme and proteinase K, refinement R-factors are much higher for MicroED structures than for X-ray structures. In our case, some of this might be improved by collecting more complete data over multiple FIB-milled crystals. However, more generally, we expect that this discrepancy will only improve as we gain a better understanding of how electrons interact with macromolecular crystals and develop data analysis software that handles processing of MicroED data and refinement of structural models based on electron scattering more appropriately.

## 5. Acknowledgements

We thank J. Rodriguez and D. Hekstra for helpful insight. FIB milling was done in the Beckman Institute Resource Center at the California Institute of Technology. M.C.T. is supported by NSF STC-1231306, a Ruth L. Kirschstein National Research Service Award (F32 HL129989), and the UCSF Program in Breakthrough Biomedical Sciences. J.S.F. is supported by a Packard Fellowship from the David and Lucile Packard Foundation, NIH GM123159, NIH GM124149, UC Office of the President Laboratory Fees Research Program LFR-17-476732, and NSF STC-1231306. N.K.S. is supported by NIH GM117126. S.I. is supported by the Platform Project for Supporting Drug Discovery and Life Science Research (Basis for Supporting Innovative Drug Discovery and Life Science Research (BINDS)) from the Japan Agency for Medical Research and Development (AMED). R.A.W. was supported by the NSF Graduate Research Fellowship. Portions of this research were carried out at the Linac Coherent Light Source (LCLS) at the SLAC National Accelerator Laboratory, supported by the DOE Office of Science, OBES under contract DE-AC02-76SF00515. The HERA system for experiments at MFX was developed by Bruce Doak and funded by the Max-Planck Institute for Medical Research. Portions of this research were performed at beamline 3 of SACLA with the approval of the Japan Synchrotron Radiation Research Institute (JASRI) (proposal no 2017B8055). We thank the staff at SACLA for their assistance. Data processing was performed in part at the National Energy Research Scientific Computing Center, supported by the DOE Office of Science, Contract No. DEAC02-05CH11231.

