## Supporting Information for "Comparing serial X-ray crystallography and microcrystal electron diffraction (MicroED) as methods for routine structure determination from small macromolecular crystals"

*†Present address: MRC laboratory of molecular biology, Cambridge, UK*

**Table of Contents:**

Supporting Figures 1-9 p. 3-8

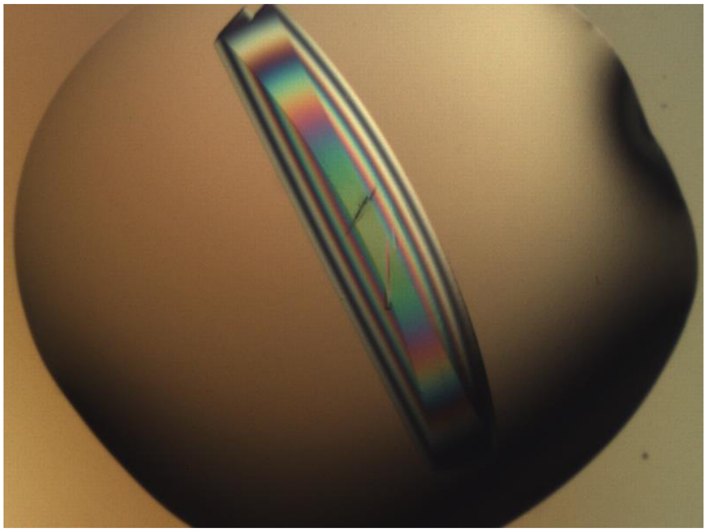

**Supporting Figure 1**. Previously, CypA crystallization conditions were optimized to produce very large single crystals. The pictured crystal (viewed using a cross-polarizer) was over 1 mm long and was visible to the naked eye.

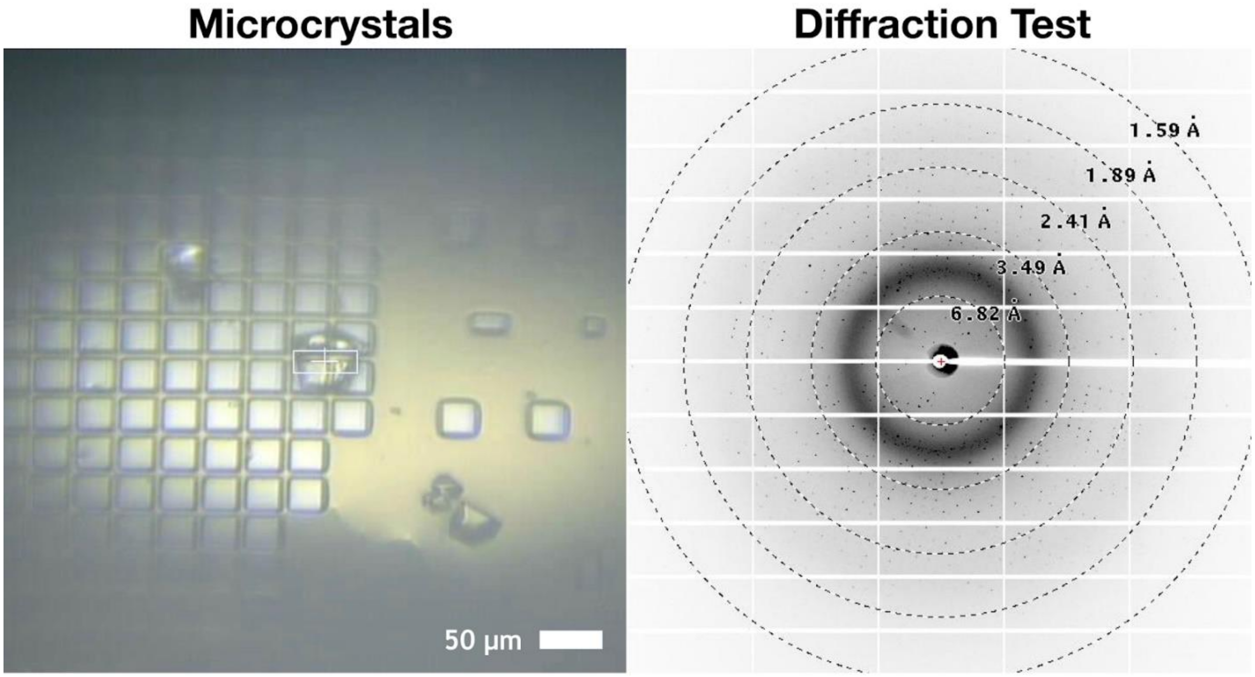


**Supporting Figure 2.** Image of microcrystals on a Mitegen micromesh grid (left) and the subsequent diffraction from one of these crystals (right). Crystals were 20-50 µm in size, and diffracted to the edge of the detector at 1.59 Å.

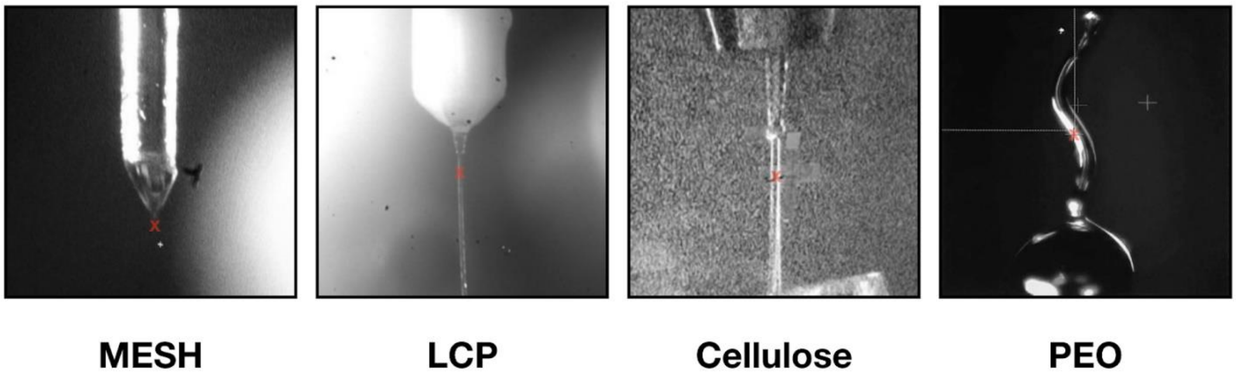


**Supporting Figure 3.** Images of jets delivering microcrystal slurries to the XFEL interaction point (red “x” in each image). Minimal viscogens were added to the crystalline slurry for the MESH injector system. When using a viscous extrusion type of injector, a variety of carrier media were tested, including LCP, Cellulose, and PEO.

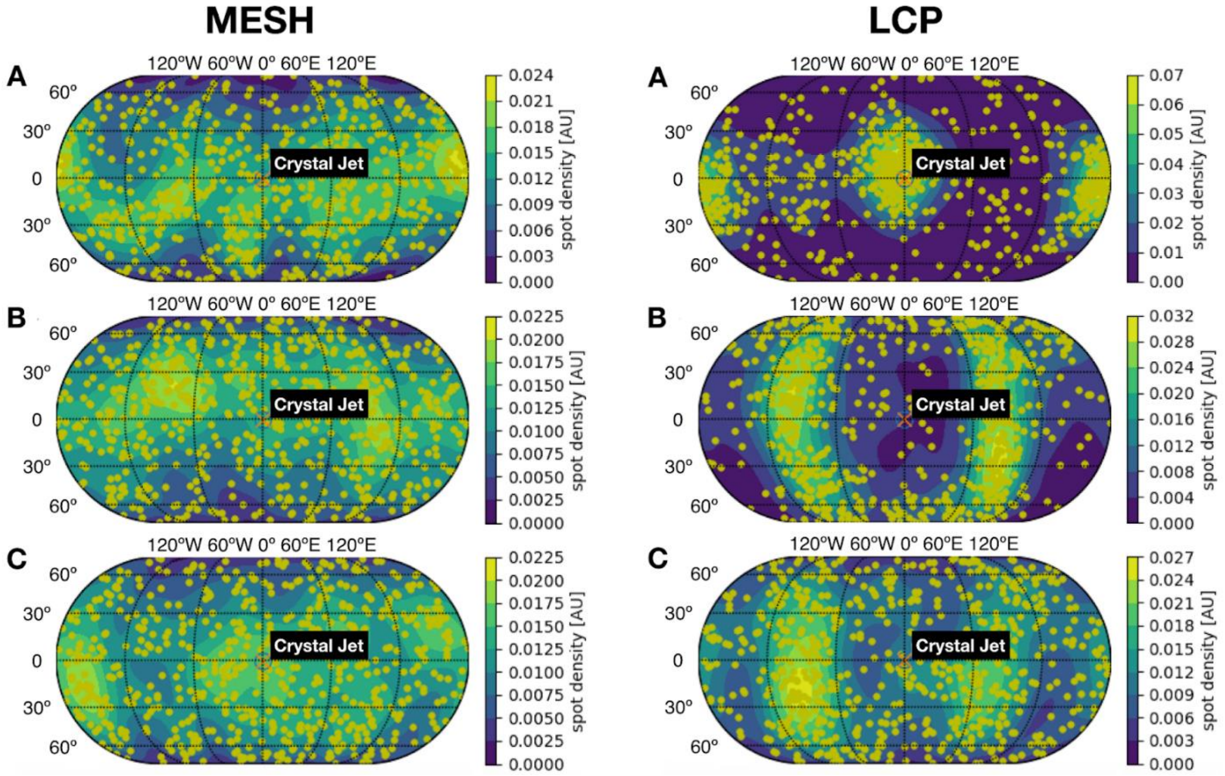


**Supporting Figure 4.** Map of crystal orientations from the MESH data collection (left panel) and from the LCP data collection (right panel). A subset of each dataset is shown for visual clarity. The orientations appear evenly distributed for the MESH data, with no major bias introduced by the electric field created by the injection system. The viscosity of the LCP carrier media appears to have induced an orientation bias, but did not prohibit collection of a complete dataset with high redundancy.

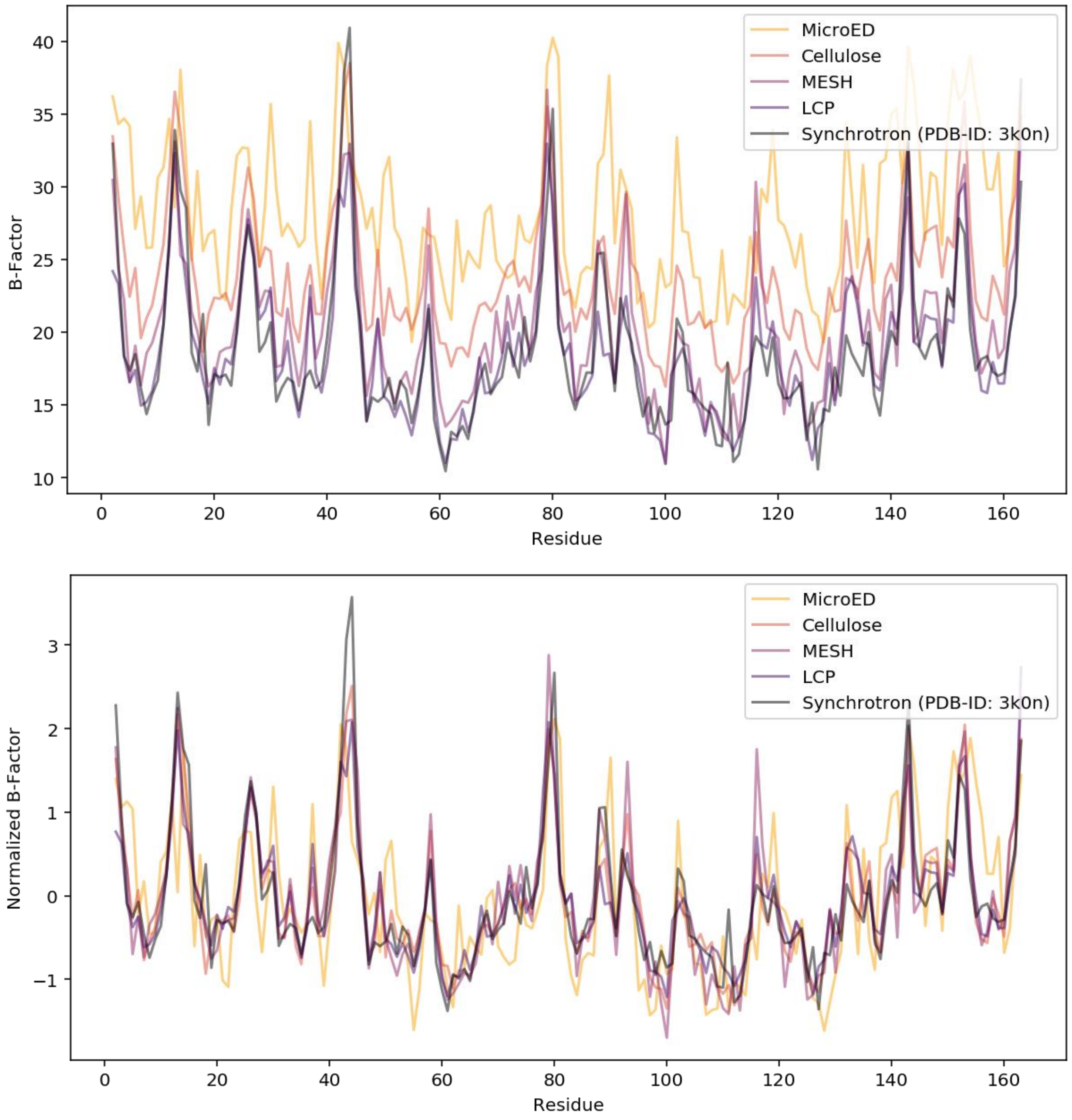


**Supporting Figure 5.** Raw (top) and normalized (bottom) B-factor per residue across data sets. A previously published structure, solved at room temperature using rotational collection from a single crystal (PDB ID *3K0N*) is provided for reference. Most variation is systematic, and thus is removed by normalization.

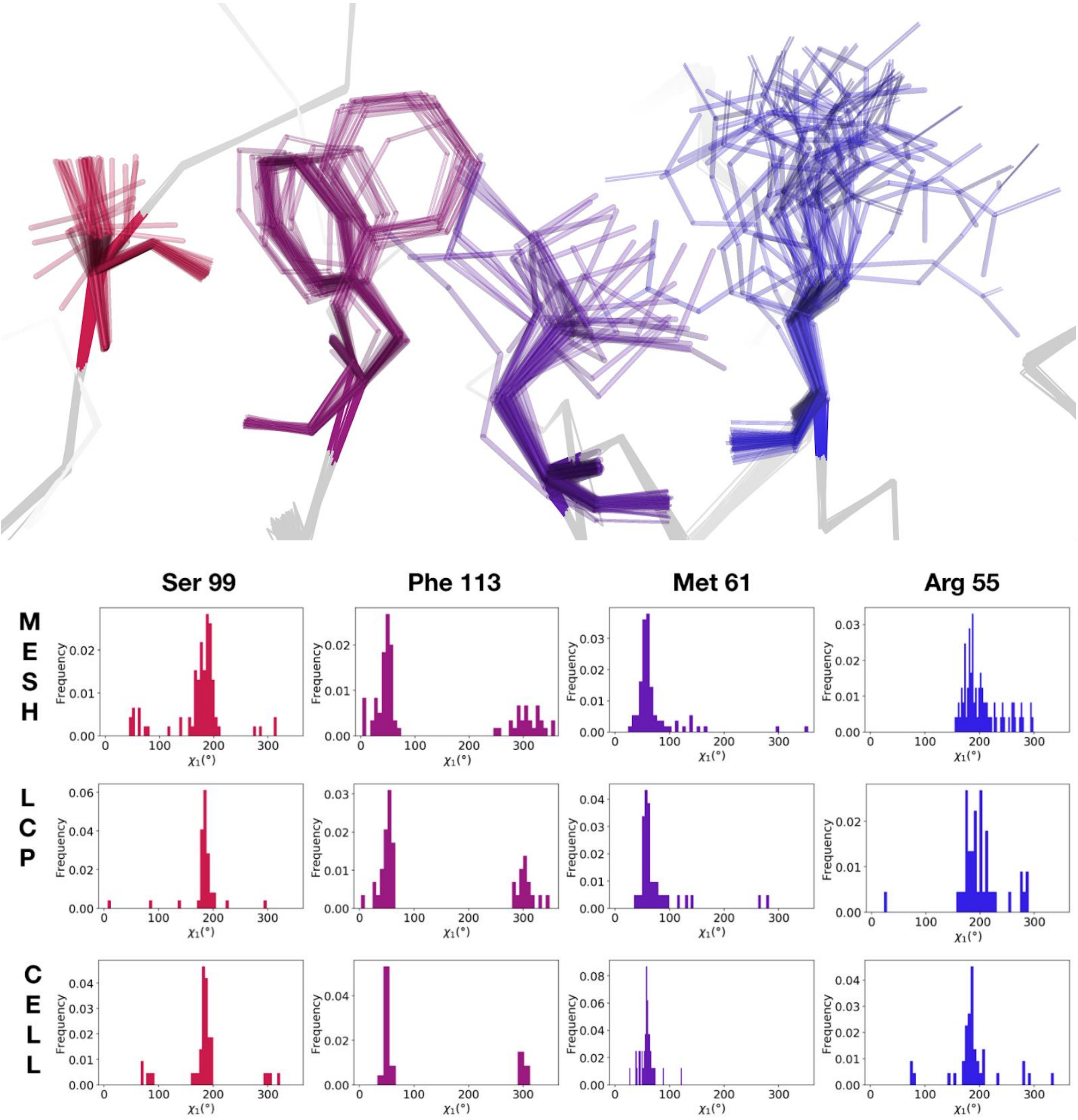


**Supporting Figure 6.** Visualization of the ensemble of conformers generated via phenix.ensemble_refine for the three serial XFEL datasets. The analysis focuses on a network of amino acid side chains that are known to be dynamic and important for function. In the top panel, the ensemble model for the cellulose dataset is displayed. Sticks are shown for the residues of interest (R55, M61, S99, F113), while the backbone is displayed as a ribbon for the rest of the structure. In the bottom panel, histograms of the distribution of chi1 angles are plotted for each of the four residues from the respective ensemble. The comparison shows that ensembles derived from the different serial X-ray experiments are essentially equivalent.

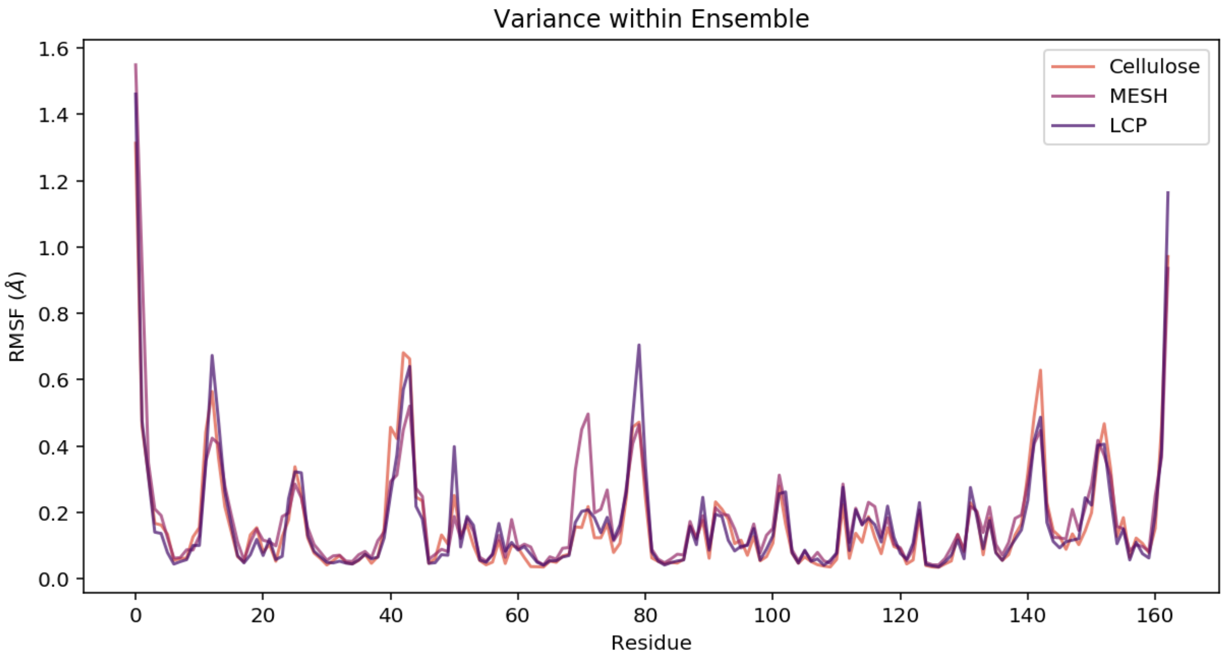


**Supporting Figure 7.** Average RMSF per residue for each ensemble generated for the serial XFEL datasets. A loop containing residues 64-74 samples more conformations in the MESH ensemble than in the other two.

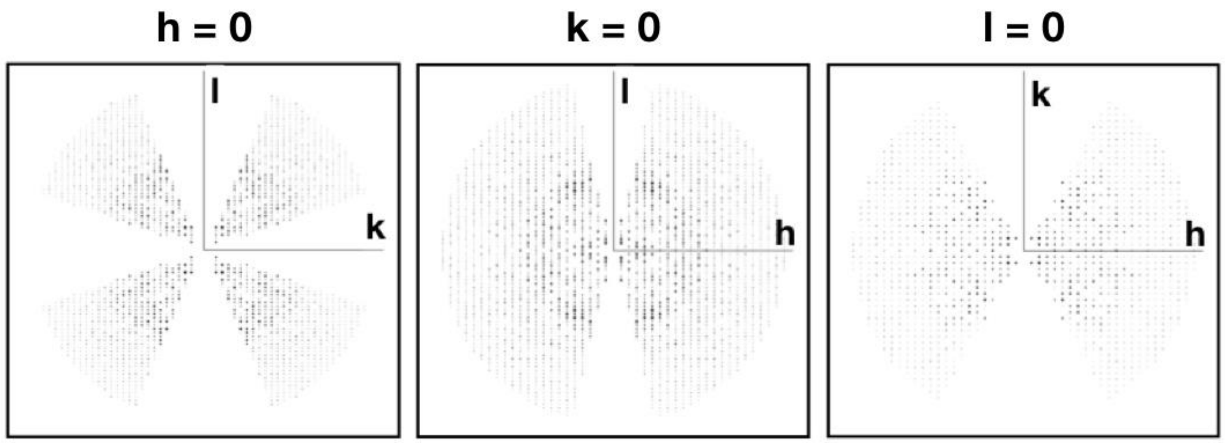


**Supporting Figure 8.** MicroED data visualized on two-dimensional slices of the reciprocal lattice. Measured reflections are visualized in black. Missing measurements along the kl plane may contribute to challenges in assignment of unit cell dimensions.

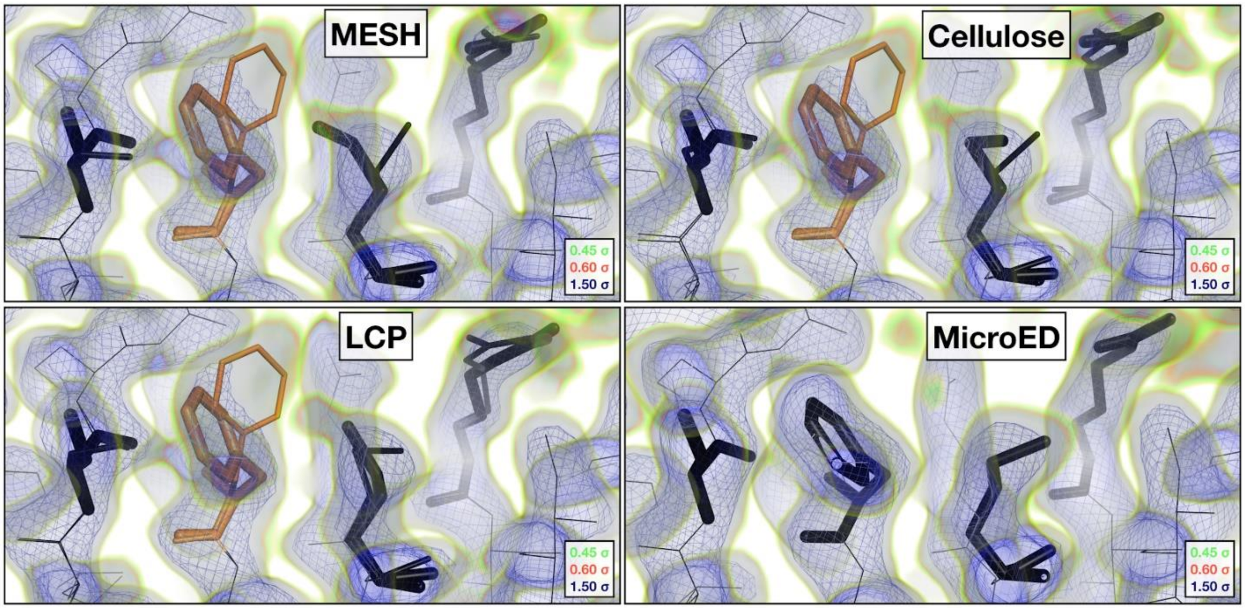


**Supporting Figure 9.** Visualization of all four datasets truncated at 2.5 Å. Comparison of the 2mFoFc maps and refined multi-conformer models reveals evidence for alternative conformations in the room temperature data (MESH, LCP, Cellulose), while cryogenic data (MicroED) supports a single conformer model.
